# Mechanistic reconstruction of receptor-to-transcription factor signaling integrating prior knowledge and omics

**DOI:** 10.64898/2026.06.19.733348

**Authors:** David Liaskos, Omid Oftadeh, Margherita Tonini, Maria Masid, Vassily Hatzimanikatis

## Abstract

Omics profiling is ubiquitous, yet resolving how signal transduction shapes cellular physiology remains challenging. Most analyses interpret omics data by summarizing catalogs of pathways or by fitting networks to observations. Here we present SIGMA, a framework that converts curated prior knowledge into causally interpretable, elementally balanced signal-transduction cascades that connect sources to targets. By enforcing elemental balance, SIGMA links processes across pathway catalogs and reveals crosstalk beyond canonical definitions. It enumerates alternative cascades to expose parallel routing and identify context-dependent essential steps. We introduce balanced cascade enrichment analysis to map omics data onto alternative cascades and rank them by context-specific molecular support. Using SIGMA, we reconstructed signal flow from receptors to transcription factors that regulate metabolism, uncovering mechanisms that control metabolic reprogramming. In CD8^+^ T cells, SIGMA identified cascades connecting TGF-β receptors to SP1 and indicated weakening of this axis in exhaustion. Overall, this framework enables mechanistic interpretation of signaling across datasets and guides the design of causal perturbations.

## Main

Advances in high-throughput technologies have enhanced our ability to describe how cells sense and transduce diverse signals, thereby providing insights into gene expression and cellular physiology^1–3^. Extensive efforts have focused on reconstructing prior knowledge signaling networks (PKNs) by consolidating pathways and biochemical evidence from years of research into databases^4,5^. These network reconstructions aim to enable systems-level interpretation of large datasets by inferring context-specific causal signal flow and thus linking inputs to cellular responses^6,7^. Achieving this goal, however, requires mechanistic reasoning beyond the statistical associations typically captured by pathway enrichment analysis using these databases^8^. It further requires overcoming reliance on *a priori* pathway definitions that underpin enrichment approaches and reflect context biases, thereby enabling inference beyond canonical boundaries and the discovery of inter-pathway interactions and crosstalk^9–11^.

Toward this goal, computational approaches have been developed to infer signal flow in networks and provide mechanistic interpretations of omics data. Steady-state formulations offer scalable mechanistic interpretation across large PKNs^12^. These methods connect candidate inputs to downstream responses by tracing paths through the network^13–15^, propagating activity across it^16,17^, localizing altered regions^18,19^, inferring dependency structures^20–24^, and extracting subnetworks that best reconcile experimental data with prior knowledge^25–29^. Dynamic formulations provide a complementary representation by modeling temporal signal flow, but they require extensive parameterization or face a combinatorial explosion of states, confining their application to small, isolated cascades^30–36^. Collectively, these methods seek to reconstruct the causal architecture of cellular responses from global network topology and omics data.

Such mechanistic approaches have yielded insights into cancer^37,38^, immune signaling^39,40^, cell-cell communication^41^, and neurodegeneration^42^, underscoring their promise for dissecting complex biological phenomena. Yet many methods still interpret data through predefined pathway catalogs, thereby carrying biases from pathway definitions into downstream analyses^8,9,43^. Most approaches return a single subnetwork or pathway summary, often by fitting networks to data, which can obscure alternative routing and increase the risk of overfitting^6,15,25–29^. Inter-pathway interactions are typically implicit, and consequently, the contribution of crosstalk is rarely assessed^12,18^. Finally, reaction mechanisms are often abstract, with limited support for process-resolved steps, such as phosphorylation-site specification and cofactor participation, constraining biochemical interpretability and the design of experimental interventions^44–47^. Together, these challenges call for an integrated mechanistic view of signaling that moves beyond pathway boundaries, captures alternative routes and crosstalk, and retains the molecular resolution needed to interpret omics data.

To unify current advances and address the remaining challenges, we present SIGMA (Signal Inference with Graph-based Mechanistic Analysis), a framework that extracts elementally balanced signaling cascades from curated networks. SIGMA enforces information-flow logic and process-level resolution, converting paths from defined sources to targets into causally interpretable cascades that recover the molecular events supporting signal transduction. It supports unbiased exploration of network-scale interconnectivity and quantifies crosstalk contributions to signal transduction by extracting cascades without fitting to a particular dataset. By exhaustively enumerating alternative cascades, SIGMA identifies essential steps shared across them and, when paired with omics data, evaluates condition-specific routing through distinct cascades. To interpret omics data, we introduce balanced cascade enrichment analysis (BCEA), which projects measurements onto extracted cascades and ranks the alternatives by their data support, providing a mechanistic surrogate for traditional pathway enrichment. Here, we apply SIGMA to delineate the receptor-to-transcription factor (TF) signaling landscape that regulates cellular metabolism. We reconstruct the network around TFs regulating metabolic genes, identify upstream receptors that converge on them, and characterize receptor-TF wiring patterns. We use the TGF-β receptor to SP1 axis to demonstrate how the framework extracts alternative balanced cascades, resolves supporting reactions, and quantifies crosstalk. Finally, using publicly available CD8^+^ T cell data, we perform BCEA to map transcriptomic profiles onto alternative cascades, identifying a transcriptionally supported TGF-β-SP1 axis in naive cells that weakens during exhaustion. These applications validate SIGMA as a framework for mechanistic explanation of signal propagation and for generating testable hypotheses about signaling-mediated metabolic reprogramming.

### The SIGMA framework

We developed SIGMA, a computational framework that converts prior knowledge into causally interpretable, elementally balanced signaling cascades and links these cascades to omics data (Fig. 1). Given a directed PKN as input, the framework starts from a set of root species, the signaling entities that define the biological question (Fig. 1a). These root species commonly include receptors, TFs, or other central proteins. SIGMA reconstructs a question-specific signaling network from the larger PKN, anchored around these roots. To do so, the framework maps the root species to their annotated pathways and expands the pathway collection along documented inter-pathway connections, adding pathway layers until further expansion yields little additional connectivity (Fig. 1a and Methods). This layered expansion defines a signaling neighborhood around the root species, ensuring broad coverage while controlling network complexity. Using this pathway collection, we perform evidence-guided gap filling to restore intra- and inter-pathway reactions supported by the selected knowledge base and the literature (Fig. 1a and Methods). Collectively, this reconstruction yields a reduced, gap-filled signaling network centered on the root species for subsequent cascade extraction.

**Fig. 1.**
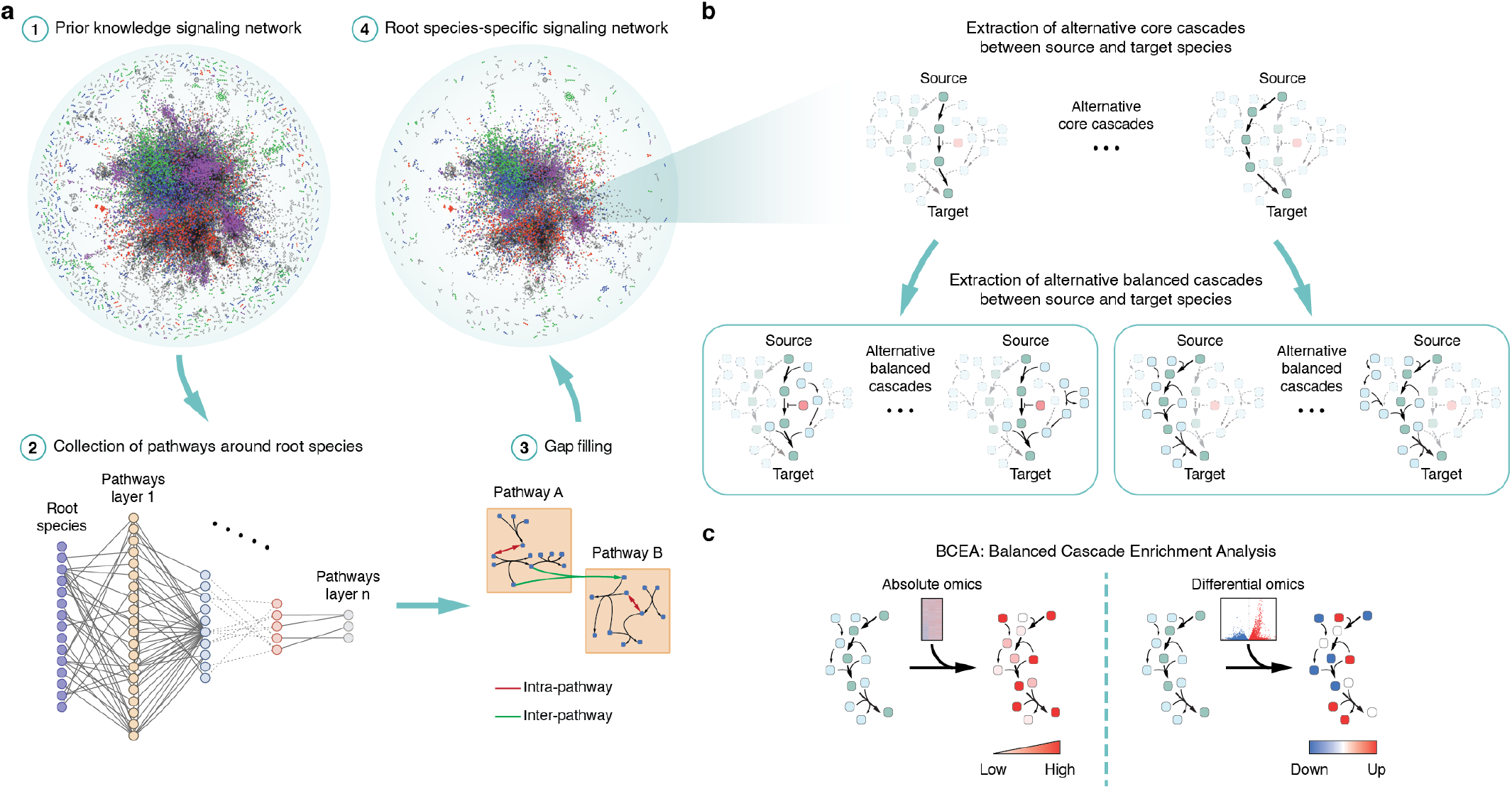
Network reconstruction, cascade extraction, and omics interpretation with the SIGMA framework. **a**, Starting from a prior knowledge signaling network, SIGMA selects root species relevant to the biological question, such as receptors or transcription factors (1). After mapping these root species to their pathways, the framework expands pathway layers along documented inter-pathway connections until added layers contribute little new connectivity (2). It then performs evidence-guided gap filling (3). The result is a reduced, root species-specific signaling network (4). Figure created with Cytoscape. **b**, From the reconstructed network, SIGMA extracts a core cascade, which is the shortest path from a selected source to a selected target. It then enumerates alternative core cascades of equal or greater length. For each core, the framework constructs an elementally balanced cascade by identifying the minimal set of additional reactions necessary to sustain information flow through the core. Multiple balanced cascades may exist per core. **c**, Balanced cascade enrichment analysis (BCEA) maps omics data onto the extracted, elementally balanced cascades. It quantifies molecular support for a cascade from absolute measurements and deregulation from differential measurements. Mapping omics data onto molecular species localizes highly expressed or deregulated regions of the cascade. SIGMA ranks alternative cascades for a given source-target pair by their BCEA enrichment scores and identifies regions whose data support shifts across alternatives.

From the reconstructed network, SIGMA first extracts a core cascade, which is the shortest directed signaling path connecting a selected source to a selected target (Fig. 1b and Methods). This core provides a parsimonious scaffold for signal traversal through the network^48^, capturing a sequence of species-to-species connections without process-level detail. The framework then enumerates alternative core cascades of the same minimal length and, where biologically relevant, longer alternatives that relax parsimony and reveal additional feasible paths. To transduce signal, a core cascade requires supporting network elements, with enabling reactions and participating species active, and inhibitory regulators inactive. For example, a core cascade may include only the state change of a protein from unphosphorylated to phosphorylated, whereas the complete phosphorylation reaction involves the kinase and ATP-to-ADP conversion. To formalize these dependencies, SIGMA extracts an elementally balanced cascade for each alternative core cascade, specifying the minimal set of additional reactions and species required to sustain transduction (Fig. 1b and Methods). The balanced cascade therefore contains the core species along with the supporting species and reactions identified during balancing. Each alternative core may yield multiple balanced cascades of the same minimal size or larger. Enumerating alternative core and balanced cascades under the parsimonious principle of minimality enables the identification of potentially essential steps shared across alternatives and cascade-specific steps that vary among them^49^. These alternatives also reveal parallel cascades that achieve the same source-target mapping and may inform condition-specific rewiring, since not all cascades transduce signal in every cellular state or cell type^50–52^. By assembling elementally balanced cascades via an unbiased network scan, SIGMA moves beyond predefined pathway annotations and canonical boundaries, combining reactions across pathways and thereby uncovering crosstalk within the transduction mechanism.

To connect the extracted cascades to omics data within SIGMA, we developed balanced cascade enrichment analysis (BCEA), which employs the complete elementally balanced cascade as the enrichment unit rather than an *a priori* defined pathway (Fig. 1c and Methods). BCEA maps transcriptomic or proteomic measurements, from bulk or single-cell datasets, onto the molecular species in each cascade, enabling mechanistic interpretation of cascade support or deregulation at the process level. Using the entire balanced cascade focuses enrichment on the specific transduction mechanism and avoids limitations of traditional pathway enrichment, which can omit required steps and crosstalk or include irrelevant pathway segments. With absolute measurements, cascade regions with a high density of expressed or abundant species indicate strong molecular support for potential signal flow, whereas regions with few such species indicate limited support. With differential measurements, BCEA localizes cascade segments with pronounced deregulation between conditions. In both measurement settings, this method can quantify the contribution of supporting reactions and crosstalk to overall cascade enrichment. Finally, SIGMA ranks alternative balanced cascades for a given source-target pair by their overall data support and pinpoints altered regions across alternatives, together indicating candidate mechanisms of condition-specific rewiring.

This framework can be applied to a plethora of biological questions, where unbiased exploration of prior mechanistic knowledge enables understanding of signal transduction across datasets.

## Results

### Signaling network reconstruction around transcription factors regulating metabolism

We implemented SIGMA to understand how intracellular signaling regulates metabolism as a downstream process. To define the transcriptional link between signaling and metabolism, we focused on TFs that regulate metabolic genes^53^. We identified 1,884 unique metabolic genes encoding metabolic enzymes from the human genome-scale model Recon3D^54^ (Methods). Using TRRUST^55^, a literature-curated resource of human transcriptional regulatory interactions that catalogs 748 TFs, we mapped direct regulatory relationships from 115 TFs to 237 of these genes (Fig. 2a, Methods and Table S1a,b). Together, these 115 TFs define signaling targets with the potential to reprogram metabolism. Although most TFs regulate only a few metabolic genes, some, including SP1, NFKB1, and HIF1A, regulate multiple and thus may influence broad regions of metabolism.

**Fig. 2.**
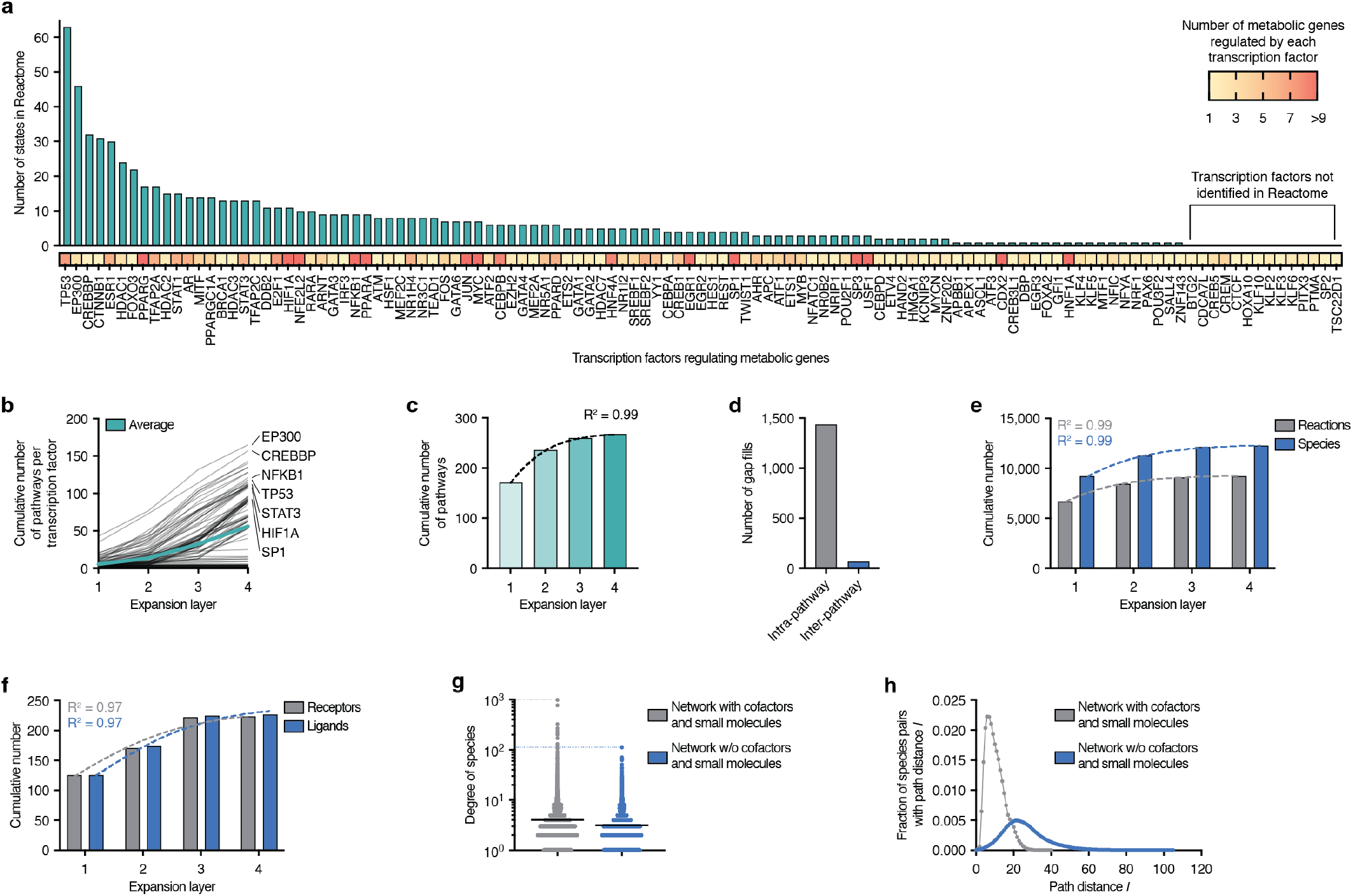
Signaling network reconstruction around transcription factors regulating metabolism. **a**, Transcription factors (TFs; n = 115) that regulate metabolic genes, with a heatmap showing the number of metabolic genes per TF. The bar plot ranks TFs by the number of Reactome states mapped per TF. Fourteen TFs lacked a corresponding Reactome species (right). **b**, Cumulative number of pathways per TF across four expansion layers. The green line shows the per-layer average, and labeled lines denote representative TFs. **c**, Collective addition of unique pathways across all TFs by expansion layer. The dashed curve shows a logistic fit (R^2^ = 0.99). **d**, Number of evidence-guided gap fills, with intra- and inter-pathway gap fills shown as separate bars. **e**, Cumulative numbers of reactions and species across four expansion layers. Dashed curves show logistic fits (both R^2^ = 0.99). **f**, Cumulative numbers of plasma membrane receptors and extracellular ligands across expansion layers. Dashed curves show logistic fits (both R^2^ = 0.97). **g**, Degree centrality distributions for species in the TF-specific network with cofactors and small molecules included (gray) and without the curated set of 54 cofactors and small molecules (blue). Dotted lines mark maxima; black solid lines mark means. **h**, Distributions of shortest-path lengths (*ℓ*) for all connected species pairs with cofactors and small molecules (gray) and without them (blue). Pairs that were not connected (*ℓ* = 0) are not shown. Connected pairs: 37,425,653 (with) and 17,867,275 (without).

To reconstruct the signaling network around TFs that regulate metabolism, we used Reactome, a signaling PKN that provides detailed process descriptions, including post-translational states, subcellular localization, and cofactor participation^56^. This process-level resolution is important for SIGMA because elemental balancing depends on molecular event detail that would be lost in networks based on protein identities alone. To capture this molecular detail, for each of the 115 TFs, we identified all states annotated in Reactome, including complexes and post-translationally modified species (for example, phosphorylated or ubiquitinated) (Fig. 2a, Methods and Table S1c). Fourteen TFs lacked a Reactome species entry, leaving 101 TFs for downstream analysis. TP53 and EP300 appeared in numerous states, reflecting both functional diversity and extensive annotation. Some TFs also appeared in heteromeric complexes, such as JUN-FOS and STAT1-STAT3 heterodimers.

Using all states of the 101 TFs as root species, we mapped them to Reactome pathways, designated as layer 1 (Extended Data Fig. 1a). EP300, CREBBP, and STAT3 mapped to multiple pathways, positioning them as integrators of diverse signals. We also observed a positive correlation between the number of TF states and the number of pathways in which a TF appeared (Extended Data Fig. 1b). To cover the broader signaling neighborhood around these TFs, we expanded the pathway collection across four layers using documented inter-pathway connectivity in Reactome (Methods). Although the pathways associated with each TF increased across all expansion layers (Fig. 2b and Table S1f), the collective addition of pathways saturated by layer four, yielding 268 pathways in total (Fig. 2c and Table S1d). This saturation indicated that four layers captured the relevant upstream and downstream context around TFs regulating metabolism. The expansion also highlighted central pathways with high documented inter-pathway connectivity, notably the PIP3-AKT signaling pathway and the RAF-MAP kinase cascade (Extended Data Fig. 1c), consistent with convergence of multiple signals linked to metabolic regulation, including control of glycolysis, the pentose phosphate pathway, and nucleotide and lipid synthesis^57,58^.

To restore continuity of signal flow across the resulting collection of 268 pathways, we performed evidence-guided gap filling by adding intra- and inter-pathway reactions supported by Reactome documentation and targeted literature checks (Fig. 2d and Methods). This step yielded the final TF-specific signaling network comprising 9,227 reactions, 12,288 species, and 6,372 genes. The cumulative numbers of reactions and species saturated at the fourth expansion layer, mirroring the pathway-level saturation (Fig. 2e and Table S1f). Within this network, we enumerated potential inputs to the TFs regulating metabolism by identifying 224 plasma membrane receptors and their extracellular ligands (Fig. 2f, Methods and Table S1e).

Cofactors and small molecules support numerous reactions and therefore act as highly connected hubs in the network. During cascade extraction, these hubs can create artificial signaling shortcuts by linking otherwise unrelated signaling species. Reactome includes such molecules, for example, ATP, water, and ions. To identify these molecules systematically, we ranked all Reactome species by degree centrality^59^ and selected 54 cofactors and small molecules by mining the highest-degree entries (Methods and Table S1h). Accordingly, to avoid biologically irrelevant cascades, we excluded these species during core cascade extraction, while retaining canonical second messengers, and later reintroduced the excluded species during elemental balancing to preserve biochemical completeness (Methods). To quantify their impact on topology, we compared the TF-specific network with and without the 54 selected species. Removing them reduced the degree of the remaining species, indicating that most proteins are linked to at least one cofactor or small molecule (Fig. 2g and Table S1i). We also measured shortest-path distances for all species pairs and found substantial path lengthening and a larger network diameter after their exclusion (Fig. 2h, Methods and Table S1i), validating the signaling shortcuts they create.

### Tracing signaling flows from receptors to transcription factors regulating metabolism

We used SIGMA to trace signal flow from each plasma membrane receptor to TFs that regulate metabolic genes. Using receptors as source species and TF states as target species, we extracted the shortest core cascade connecting each receptor-TF state pair in the TF-specific network (Methods). These connections reflect the annotation space of the databases used in this study. Nearly half of receptors connected to at least one TF, indicating broad potential for receptor-driven metabolic regulation (Extended Data Fig. 2a and Table S2a). Linked receptors converged on a shared TF set, with TGF-β and IFN-γ receptors reaching the largest fraction of TFs. Notably, only the TGF-β and IFN-γ receptors connected to SP1, the TF with the most metabolic gene targets, suggesting a targeted route through which immune and morphogen cues may access extensive metabolic control. From the TF perspective, 77 of the 101 TFs were reachable from at least one receptor, while most of these linked TFs were reached by a common set of receptors (Extended Data Fig. 2a and Table S2a). STAT1 and STAT3 were the most broadly reachable TFs, with exclusive links to some cytokine receptors, including the IL-18 and IL-9 receptors.

To identify patterns of signaling toward TFs regulating metabolism, we examined how receptor families contributed to TF reachability (Methods). Cytokine receptors had the largest share of receptor-TF links, followed by receptor tyrosine kinases (Fig. 3a and Table S2b). By contrast, although integrins were abundant, they contributed few links, and some families, such as scavenger receptors, had none, suggesting that these inputs were unlikely to directly control metabolic programs in the annotated network. To identify the most direct routes among receptor families, we compared their path lengths to linked TFs (Methods). TGF-β receptors showed the shortest paths, whereas the death-receptor family exhibited the longest (Fig. 3b and Table S2c). Clustering of families by path-length profiles separated cytokine receptors from receptor tyrosine kinases, consistent with their different modes of access to metabolism^60,61^ (Extended Data Fig. 2b and Supplementary Fig. 1b). Together, these results position cytokine and growth factor inputs as major contributors to downstream metabolic regulation. We also examined TF families to assess which regulatory programs were most connected to surface sensing (Methods). TFs annotated for RNA polymerase II transcription and cellular stress responses were most linked to receptors (Extended Data Fig. 2c and Table S2d). In contrast, some TF families, such as RNA polymerase III transcription, lacked receptor links, indicating no direct receptor-driven control of those families in this analysis.

**Fig. 3.**
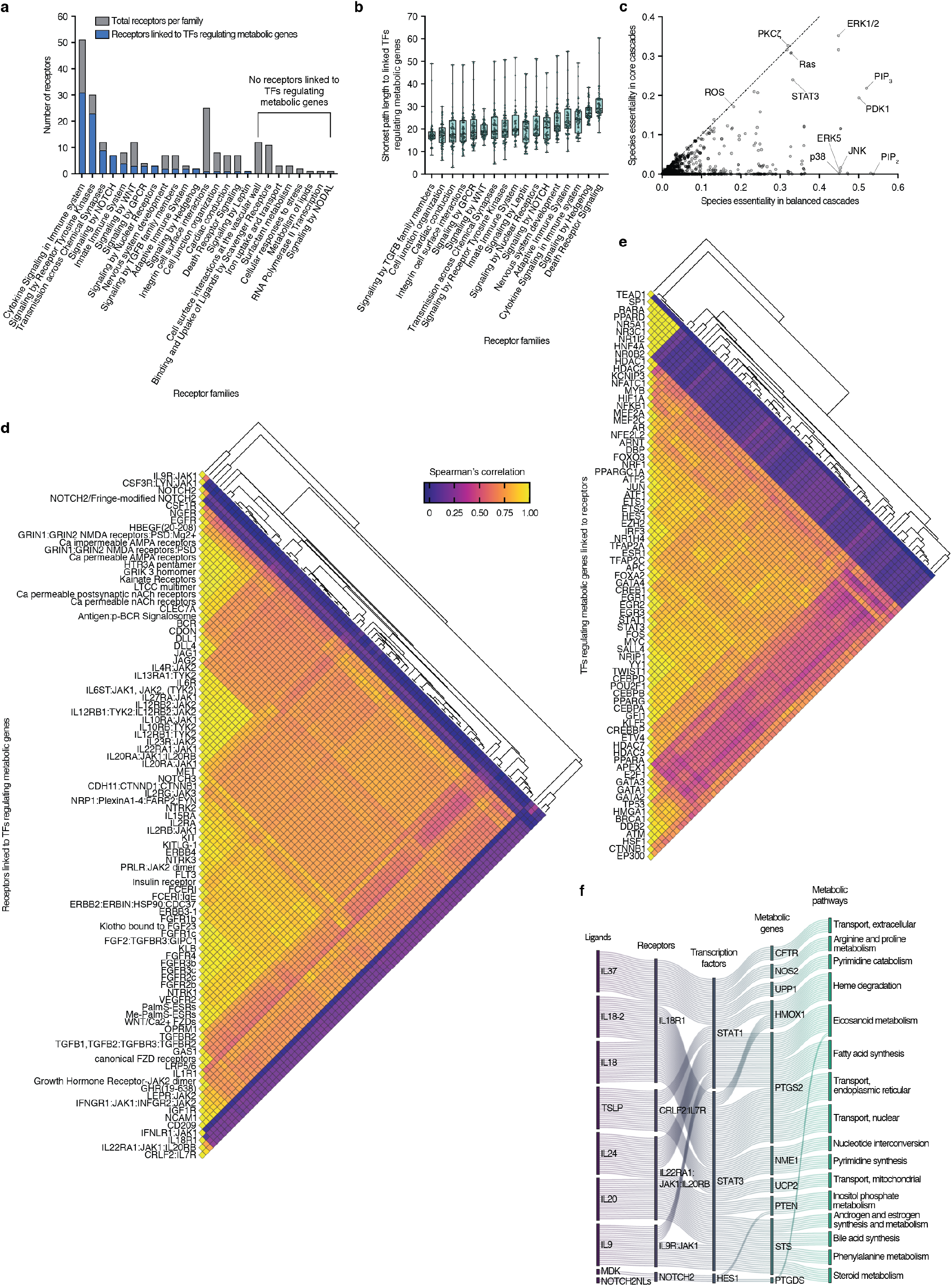
Topology and properties of information flows from receptors to metabolism. **a**, Number of plasma membrane receptors per receptor family in the TF-specific network (gray) and number linked to TFs that regulate metabolic genes (blue). Bars are ordered by the count of linked receptors. Eight receptor families had no linked receptors (right). **b**, Box plots of shortest-path lengths from receptors to linked TFs regulating metabolic genes, grouped by receptor family and ordered by the median per family. Each box spans the first to third quartiles with the median as a horizontal line. Whiskers indicate the minimum and maximum. Each point corresponds to a single TF and marks the mean shortest-path length from all receptors in that family to that TF. **c**, Global species essentiality across weighted subnetworks upstream of linked TFs. Balanced essentiality (x-axis) and core essentiality (y-axis) reflect the participation frequency of each species across TF-upstream subnetworks reconstructed from balanced and core alternatives, respectively (Methods). Each point represents one signaling species (n = 3,748); 1,512 species had nonzero core essentiality, whereas the remaining 2,236 species appeared only in balanced cascades. Highly essential species are labeled. The dashed line shows the identity line (x = y), indicating equal core and balanced essentiality. **d**, Heatmap of Spearman’s correlations among balanced downstream subnetworks of receptors linked to TFs that regulate metabolic genes (n = 93), with hierarchical clustering (Spearman distance, average linkage). **e**, Heatmap of Spearman’s correlations among balanced upstream subnetworks of TFs that regulate metabolic genes and are linked to receptors (n = 77), with hierarchical clustering (Spearman distance, average linkage). **f**, Sankey diagram illustrating how extracellular cues regulate metabolism. It shows information flow from extracellular ligands that bind plasma membrane receptors, through transcription factors that regulate metabolic genes, to metabolic pathways that control metabolic flux. The top five receptors, selected by median shortest-path length, are displayed.

Shortest-path analysis recovered the receptor-TF connectivity roadmap and provided a proxy for signaling directness. Yet it did not identify which biochemical intermediates are essential, nor the dependencies sustaining signal transduction. To move beyond simple connectivity, we used a two-step enumeration (Methods). For each receptor-TF state pair, we first enumerated all shortest-length alternative core cascades. For each alternative core, we then enumerated all minimal-size balanced cascades. For each TF, we aggregated the core and balanced alternatives separately into weighted subnetworks upstream of the TF (Methods). From these subnetworks, we computed global species essentiality, defined as the participation frequency of each species across TF-upstream subnetworks (Methods). This essentiality measure quantified how broadly and consistently each species supported receptor-mediated reachability of TFs. Comparing core and balanced subnetworks showed that core cascades underestimated the essentiality of many species and missed dependencies that emerged only after balancing (Fig. 3c and Table S2e). ERK1/2, PKCζ, Ras, and STAT3 remained among the most essential species in both representations. By contrast, PIP3 and PDK1 exceeded an essentiality of 0.5 after balancing, more than twice their core values. PIP2 had the highest balanced essentiality, reflecting its recurrent participation as a supporting dependency that appeared only in balanced cascades. Similarly, the MAP kinases JNK, ERK5, and p38 showed high essentiality only after balancing. These results show that balanced cascades expose non-core biochemical dependencies that sustain reachability to TFs and nominate multiple MAP kinases and lipid messengers, among other signaling hubs, as candidate leverage points for metabolic gene programs^57,58^.

Balanced subnetworks reveal which receptors are likely to generate similar downstream signals and which TFs are likely to receive similar upstream signals. In addition to the subnetworks upstream of each TF, we built weighted subnetworks downstream of each receptor by aggregating all minimal balanced alternatives from that receptor to its linked TFs (Methods). We computed pairwise correlations among balanced receptor-downstream subnetworks and among balanced TF-upstream subnetworks using species participation weights as variables (Methods). Higher correlations indicated receptors or TFs with structurally more similar balanced signaling subnetworks. For receptors, major clusters emerged, with the largest comprising interleukin receptors, including IL-6, IL-10, IL-12, and IL-20 (Fig. 3d, Supplementary Fig. 3a and Table S2f). A second cluster captured an ion-channel receptor module, suggesting calcium-dependent signal transduction. A third cluster grouped fibroblast growth factor receptors, revealing a canonical receptor tyrosine kinase axis into metabolic control. A few receptors, including some cytokine receptors and NOTCH2, showed near-zero correlation with all others, indicating unique downstream signatures and matching their specificity in the receptor-TF roadmap. On the TF side, the same analysis revealed a large cluster that included SP1 and nuclear receptors such as RARA and HNF4A (Fig. 3e, Supplementary Fig. 3b and Table S2g). This cluster was reachable only from TGF-β and IFN-γ receptors and was uncorrelated with other TFs, positioning these receptors as specialized entry points to metabolic reprogramming. Another cluster grouped SALL4, YY1, and TWIST1, suggesting common upstream control of cell-state programs. Smaller modules grouped related TFs, including ATF/JUN and STAT1/STAT3, consistent with shared signaling inputs.

We used the receptor-TF roadmap to trace information flow from extracellular cues to specific metabolic pathways. We selected the five receptors with the shortest distances to TFs (Supplementary Fig. 1a and Methods), including interleukin receptors and NOTCH2. We followed information flow from ligand to receptor, from receptor to TF, and from TF to metabolic gene targets and the pathways those genes control (Fig. 3f). Interleukin receptors signal to STAT1 and STAT3^60^, whereas NOTCH2 signals to HES1^62^. Downstream, STAT1/3 activate HMOX1 among their target genes, linking cytokine stimulation to the heme degradation metabolic pathway^63^. Additional targets implicated fatty acid synthesis, nucleotide interconversion, and inositol phosphate metabolism.

### Resolving TGF-β signaling and crosstalk toward SP1-mediated metabolic regulation

The receptor-TF map positioned TGF-β receptors among the most broadly connected and direct inputs to TFs regulating metabolism because they reached the largest fraction of linked TFs through comparatively short paths (Fig. 3b and Extended Data Fig. 2a). At the transcriptional-output level, our mapping of TFs to metabolic genes nominated SP1 as a prominent downstream effector with potentially broad influence on metabolism because it regulates the largest number of metabolic genes among the mapped TFs (47 genes; Fig. 2a, heatmap and Table S1c). Guided by these observations and prior evidence that TGF-β signaling engages SP1^64–66^, we used SIGMA to extract and interrogate the cascades that connect TGF-β receptors to SP1 within the annotation space of the databases used.

We defined a reference scaffold for resolving TGF-β to SP1 signal transduction by extracting the shortest core cascade from TGF-β receptor 2 (TGFBR2) to the nuclear complex of SP1 with the SMAD heterotrimer, the only SP1 state reachable from any receptor in the network (Methods). The core comprised nine steps and captured the canonical flow from ligand engagement to SP1 activation^64^ (Fig. 4a and Supplementary Fig. 4a). Briefly, dimeric TGF-β1 ligand binds TGFBR2, which recruits and phosphorylates TGF-β receptor 1 (TGFBR1). The activated receptor complex internalizes into early endosomes, recruiting SARA (ZFYVE9) and R-SMADs, SMAD2 and/or SMAD3. Activated TGFBR1 then phosphorylates the SMAD2/3 dimer, which dissociates from the receptor complex and binds SMAD4 in the cytosol. The SMAD2/3:SMAD4 heterotrimer translocates to the nucleus, where it binds SP1 and enhances its DNA binding and transcriptional activity^65,66^.

**Fig. 4.**
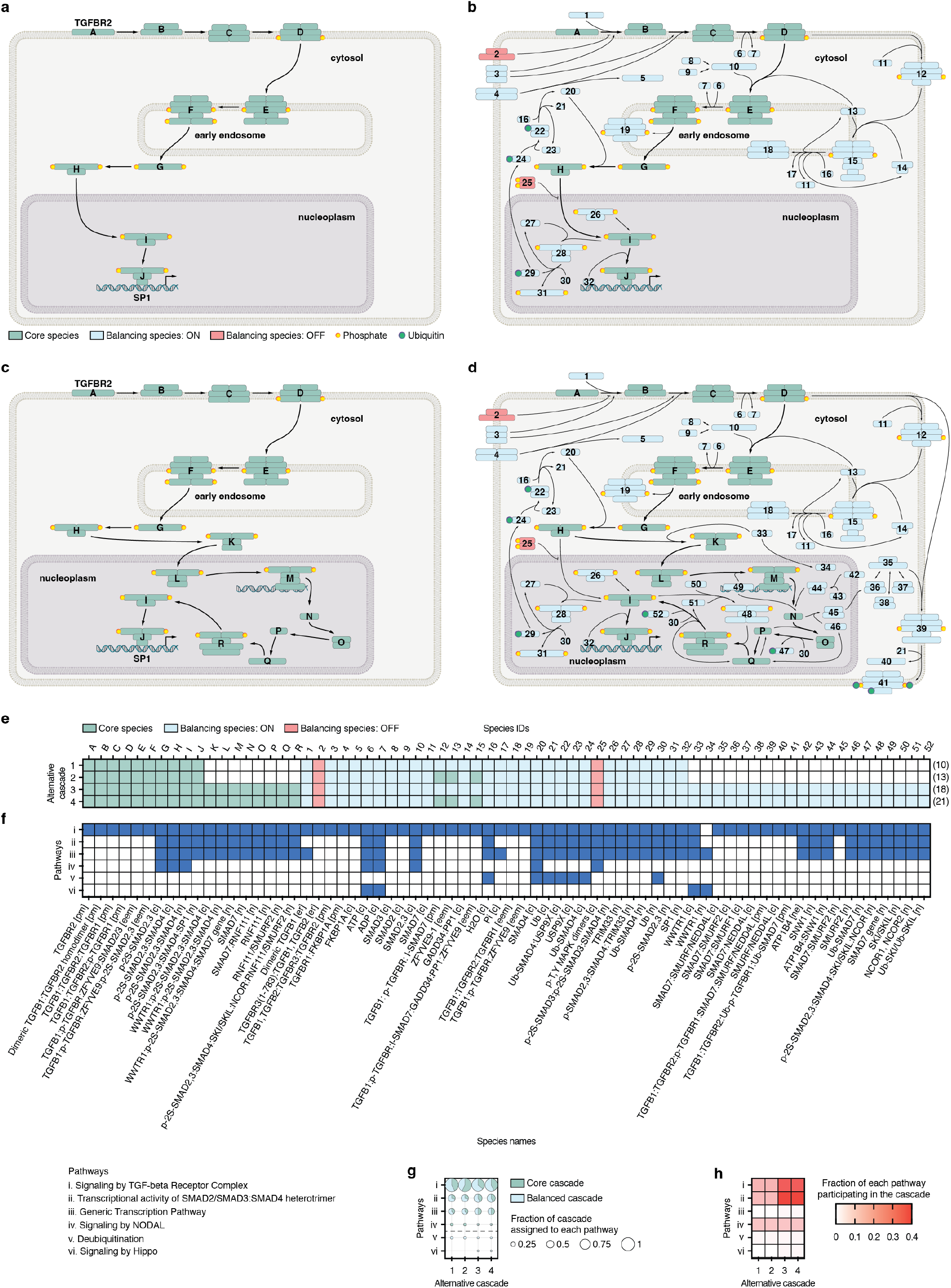
Mechanistic cascades from TGF-β to SP1 with pathway-level context. **a**, Nine-step shortest core cascade from TGFBR2 (ID A) to the nuclear complex of SP1 with the SMAD2/3:SMAD4 heterotrimer (ID J). Directed edges indicate the order of signaling events. Species IDs (for example, A and J) refer to schematic labels defined in panels **e** and **f** and are also used in subsequent panels. **b**, Elementally balanced cascade for the nine-step core in panel **a**, assembled with the minimal supporting reactions and species (23 reactions; 42 species). **c**, Alternative seventeen-step core cascade from TGFBR2 to the nuclear SP1 complex that branches via WWTR1 (ID K). Directed edges indicate the order of signaling events. **d**, Elementally balanced cascade for the seventeen-step core in panel **c** (51 reactions; 70 species). Additional reactions and species are present owing to inclusion of the WWTR1-dependent branch relative to the balanced nine-step core in panel **b**. Compartments (cytosol, early endosome, nucleoplasm) are indicated; core species use lettered IDs (green); balancing species use numeric IDs (blue); inhibitor OFF states are shown in red; post-translational modifications (phosphorylation, ubiquitination) are marked where present (**a-d**). Expanded views corresponding to panels **a-d** are shown in Supplementary Figs. 4a,b and 6a,b, respectively. **e**, Summary map of species participation across alternative cascades. Columns list species IDs. The size of each core alternative (number of species) is given in parentheses. **f**, Mapping of species to pathway annotations. Columns list species names aligned to the IDs in panel **e**. Colored boxes indicate Reactome pathway membership. In species names: pm, plasma membrane; eem, early endosome membrane; c, cytosol; n, nucleoplasm; er, extracellular region; ne, nuclear envelope. **g**, Dot plot showing, for each alternative cascade, the fraction of the cascade assigned to each Reactome pathway (i-vi, as in panel **f**) based on reaction counts. Each bubble contains a pie chart indicating the subfraction present in the core (green) versus added by balancing (blue). Pathways plotted below the dashed line were identified only after balancing. **h**, Heatmap showing, for each alternative cascade, the fraction of each Reactome pathway (i-vi, as in panel **f**) covered by the balanced cascade based on reaction counts.

To capture mechanisms missing from the core cascade, SIGMA constructed an elementally balanced cascade with the minimal supporting reactions and species (Methods). This analysis yielded a unique balanced cascade comprising 23 reactions, including six introduced by curated intra-pathway gap fills (Fig. 4b and Supplementary Fig. 4b). Balancing recovered active supporting mechanisms, including a nuclear-cytosolic SMAD4 recycling module via ubiquitination-deubiquitination. TRIM33 monoubiquitinates SMAD4 in the nucleus, while USP9X deubiquitinates it in the cytosol, sustaining reassembly of the SMAD2/3:SMAD4 heterotrimer^67^. Balancing also resolved that recruitment of SARA engages a SMAD7-dependent feedback module that, together with PP1 and GADD34, promotes TGFBR1 dephosphorylation and dampens downstream signaling^68–70^. Context-specific data can determine whether this feedback dominates over the phosphorylation branch. In addition to these supporting mechanisms, balancing identified two inhibitory species that should remain inactive for signal propagation. These were the soluble TGF-β receptor 3 (TGFBR3), which inhibits TGF-β1 binding to TGFBR2^71^, and the phosphorylated ERK1/2 dimer, which negatively regulates nuclear translocation of the SMAD2/3:SMAD4 heterotrimer^72^. The balanced cascade also quantified the energetic cost of sustaining stoichiometric signal flow. It required 14 ATP, with 12 for TGFBR1 transphosphorylation by TGFBR2 and two for R-SMAD phosphorylation (Supplementary Fig. 4b and Methods).

We exhaustively enumerated alternative core cascades between TGFBR2 and the SP1 complex and balanced each (Methods). Four cores emerged, including the nine-step core described above and longer cores of 12, 17, and 20 steps, each yielding a single balanced cascade. The twelve-step core added a receptor-proximal segment in which SMAD7 recruits PP1 via GADD34 at the activated TGF-β receptor complex to enforce SARA recruitment^68–70^ (Supplementary Fig. 5a). Because this module was already required to balance the nine-step core, the 9- and 12-step cores converged to the same balanced cascade, indicating a single mechanistic cascade rather than distinct ones (Supplementary Fig. 5b). This convergence suggests that alternative signaling paths reported in databases and the literature can rely on common enabling mechanisms. The seventeen-step core diverged at nuclear entry, where the phosphorylated SMAD2/3:SMAD4 heterotrimer first binds WWTR1 and then translocates to the nucleus to induce SMAD7 transcription^73^ (Fig. 4c and Supplementary Fig. 6a). Induced SMAD7 engages RNF111 and SMURF2 with SKI/SKIL to attenuate SMAD activity before the cascade reconverges on the nuclear SMAD2/3:SMAD4 heterotrimer that binds SP1^74,75^. The twenty-step core followed the same WWTR1-dependent cascade and added the SMAD7-PP1-GADD34 segment proximal to the receptor, as in the twelve-step core (Supplementary Fig. 7a). After balancing, the 17- and 20-step cores yielded the same enabling set of 51 reactions, mirroring the convergence of the two shorter cores into one mechanistic cascade (Fig. 4d, Supplementary Fig. 6b and Supplementary Fig. 7b).

**Fig. 5.**
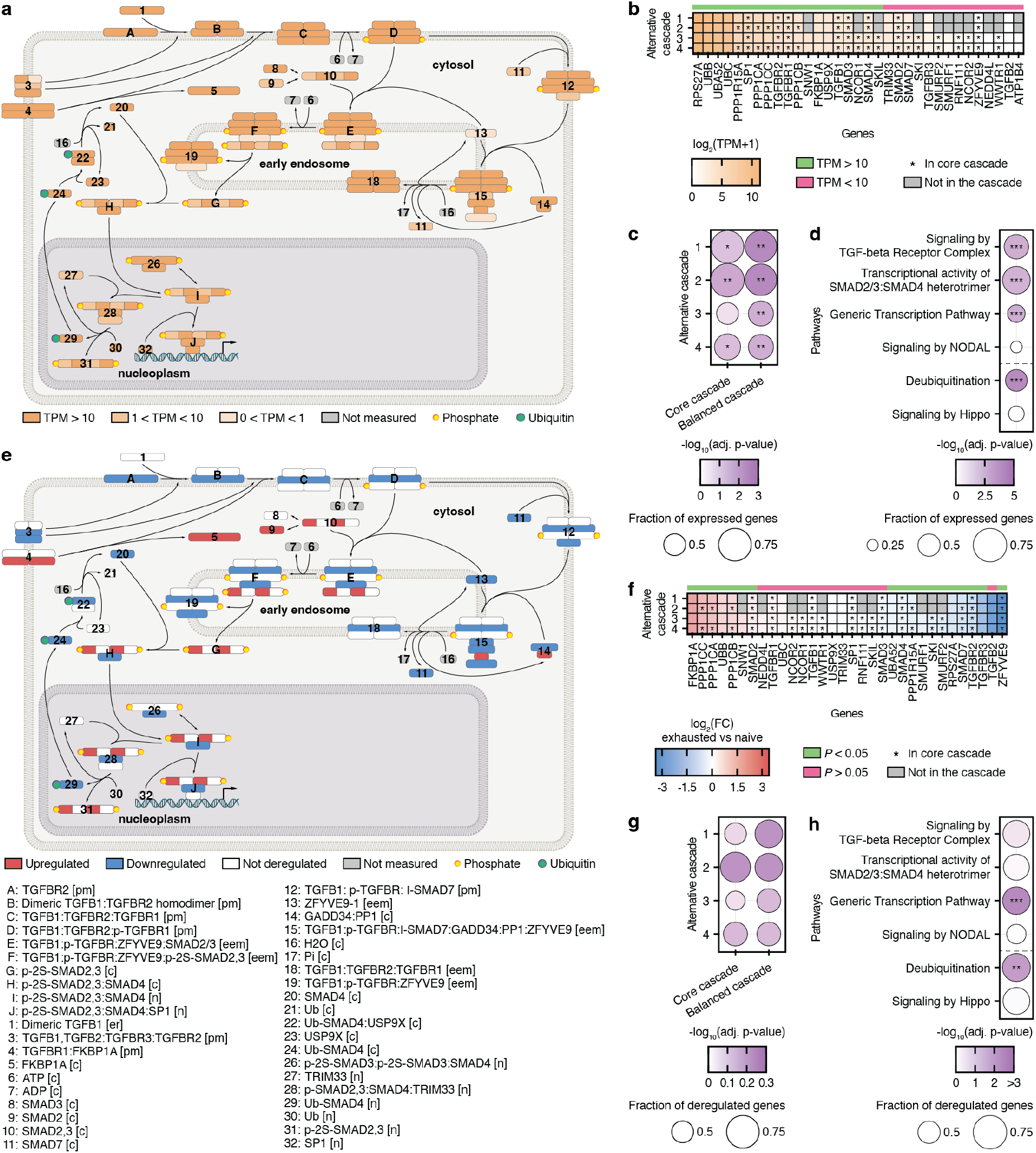
Enrichment of the TGF-β to SP1 cascade in naive and exhausted CD8^+^ T cells. **a**, Mapping of absolute expression from naive CD8^+^ T cells onto the balanced cascade shared by the 9- and 12-step cores. Color intensity (orange) indicates transcripts per million (TPM) for each UniProt-mapped gene represented in the cascade species (Methods). Protein-complex species show separate color intensities for the UniProt-mapped gene associated with each constituent protein. **b**, Heatmap showing gene expression in naive CD8^+^ T cells for all genes appearing across the alternative cascades. TPM ≥ 10 (top band) marks highly expressed genes. **c**, Dot plot of enrichment (adjusted p-value) for the core and balanced cascades of each alternative, computed from absolute expression. Bubble size indicates the fraction of highly expressed genes (TPM ≥ 10) within the cascade. **d**, Dot plot showing the enrichment (adjusted p-value) of Reactome pathway annotations participating in the alternative cascades, calculated from absolute expression. Bubble size represents the fraction of highly expressed genes (TPM ≥ 10) within the pathway. Pathways plotted below the dashed line were identified only after balancing. **e**, Mapping of differential expression between exhausted and naive CD8^+^ T cells onto the balanced cascade shared by the 9- and 12-step cores. Color indicates differential expression status for each UniProt-mapped gene represented in the cascade species (Methods). Protein-complex species show separate colors for the UniProt-mapped gene associated with each constituent protein. Upregulated genes (red): log_2_(fold change) ≥ log_2_(1.3) and adjusted *P* ≤ 0.05; downregulated genes (blue): log_2_(fold change) ≤ −log_2_(1.3) and adjusted *P* ≤ 0.05; others are not deregulated (white). **f**, Heatmap representation of fold changes (exhausted vs naive) for genes belonging to the alternative cascades. Adjusted *P* values (top band) indicate significance of differential expression and were calculated with DESeq2 (Benjamini-Hochberg corrected). **g**, Dot plot of enrichment (adjusted p-value) of both the core and balanced cascades of each alternative, calculated from differential expression. Each bubble indicates the fraction of significantly deregulated genes (|log_2_(fold change)| ≥ log_2_(1.3) and adjusted *P* ≤ 0.05) within the cascade. **h**, Dot plot of enrichment (adjusted p-value) for the Reactome pathway annotations participating in the alternative cascades, computed from differential expression. Bubble size shows the fraction of significantly deregulated genes (|log_2_(fold change)| ≥ log_2_(1.3) and adjusted *P* ≤ 0.05) within the pathway. Enrichment was performed at the gene level with over-representation analysis using the upper tail of the hypergeometric cumulative distribution and Benjamini-Hochberg correction (**c**,**d**,**g**,**h**); asterisks denote significance (*adjusted p-value ≤ 0.05, **adjusted p-value ≤ 0.01, ***adjusted p-value ≤ 0.001).

Because enumeration produced multiple feasible cascades, we used their intersection to nominate essential mechanisms for this source-target pair (Methods). All species required to balance the 9- and 12-step cores also appeared in the longer balanced alternatives, identifying them as essential for TGF-β to SP1 transduction (Fig. 4e and Table S3a). Applying the same logic at the reaction level showed that all reactions required for the shorter balanced cascades were shared across all alternatives, whereas additional reactions introduced by longer alternatives were cascade-specific (Supplementary Fig. 8a and Table S3b). Together, the species and reaction essentiality patterns highlighted receptor complex activation, SMAD regulation, and SMAD4 recycling as foundational elements that maintain signal integrity into SP1. These nominations provide a compact set of perturbation points expected to modulate TGF-β access to SP1 regardless of cascade.

To determine how these mechanisms were distributed across pathway annotations, we mapped cascade reactions to Reactome pathways (Methods). Signaling by the TGF-β receptor complex accounted for the largest fraction of every alternative (>85%; Fig. 4f,g, Supplementary Fig. 8b and Table S3e-g). Transcriptional activity of the SMAD2/3:SMAD4 heterotrimer and the generic transcription pathway also accounted for substantial cascade fractions, approximately 45% and 35%, respectively. However, these annotations overlapped extensively because the SMAD transcriptional program is nested within both TGF-β receptor signaling and generic transcription^56^. Accordingly, cascade segments were covered by multiple annotations simultaneously (Fig. 4f, Supplementary Fig. 8b and Table S3e,f). Pathway mapping also revealed potential crosstalk. Signaling by Nodal mapped to SMAD heterotrimer formation and translocation, consistent with crosstalk via phosphorylation of R-SMADs by Nodal receptors^76,77^. Deubiquitination appeared only after balancing, reflecting the USP9X-dependent SMAD4 recycling module^67^. Hippo signaling appeared only in the longer balanced alternatives, describing WWTR1-dependent crosstalk rather than an obligate module^73,77^.

Finally, we quantified the fraction of each pathway annotation assigned to each alternative balanced cascade to identify which pathways were more likely to modulate the TGF-β-SP1 axis (Methods). Signaling by the TGF-β receptor complex and transcriptional activity of the SMAD2/3:SMAD4 heterotrimer showed the largest internal coverage, followed by Nodal signaling (Fig. 4h and Table S3h). This result reinforced Nodal signaling as a plausible crosstalk mechanism. Other pathway annotations contributed only small fractions, indicating that only specific segments were involved rather than broad pathway programs.

### Recovering a transcriptionally supported TGF-β-SP1 axis in naive CD8^+^ T cells that weakens in exhaustion

TGF-β signaling is a central regulator of adaptive immunity and a driver of CD8^+^ T cell exhaustion, reducing cytotoxic function and limiting control of tumors and chronic infections^78,79^. Here, we used BCEA under the SIGMA framework to ask whether naive CD8^+^ T cells provide molecular support for potential signal transduction from TGF-β receptors to SP1 and whether exhaustion rewires this axis, with implications for metabolic reprogramming.

We mapped absolute transcriptomic profiles of naive CD8^+^ T cells^80^ (Extended Data Fig. 3a,b, Methods and Table S4a,b) onto the balanced cascade connecting TGF-β to SP1 to contextualize the cascade and assess its transcriptional support. The balanced cascade shared by the 9- and 12-step cores was highly supported, with 16 of 22 genes highly expressed and four expressed at moderate levels (Fig. 5a,b, Extended Data Fig. 3d, Methods and Table S4c). These genes included the receptor pair TGFBR1/2, the core transducers SMAD2/3/4, and SP1. The deubiquitinase USP9X and ligase TRIM33 showed robust expression consistent with a SMAD4 ubiquitination-deubiquitination cycle. In contrast, the endosomal recruiter SARA (ZFYVE9) was expressed at low levels and TGFB2 was nearly absent, pointing to receptor-endosome assembly as a potential limiting step. SMAD7 expression was lower than that of other SMADs, implying that inhibition at the receptor complex was less prominent than activation. By comparison, the balanced cascade shared by the two longer cores showed weaker transcriptional support, with only three of the eleven additional genes highly expressed (Fig. 5b, Extended Data Fig. 3c,d, Methods and Table S4c). WWTR1 was nearly absent, while its downstream components, including RNF111, SMURF2, SKI, and SMAD7, were expressed at low to moderate levels, indicating that Hippo-mediated crosstalk was unlikely in naive cells. ATP1B4 was undetected and, together with relatively low SMAD7, indicated a limited inhibition module. Overall, these results suggested that naive cells favor the shorter cascade, since modules unique to the longer alternatives showed limited transcriptional support.

We used BCEA to quantify these trends, demonstrating that the balanced cascade from the two shorter cores had higher enrichment than the longer pair (Fig. 5c, Methods and Table S4d). Enrichment on the cores alone was consistently lower than on their balanced counterparts, indicating that supporting modules were transcriptionally present. Thus, supporting mechanisms may provide candidate points for modulating TGF-β to SP1 transduction beyond the core. Within the TF-specific network, TGF-β receptor complex signaling, transcriptional activity of the SMAD2/3:SMAD4 heterotrimer, and generic transcription were among the most enriched pathways (Fig. 5d, Extended Data Fig. 3e, Methods and Table S4e). This analysis highlighted the importance of elemental balancing, as deubiquitination, which emerged only after balancing, was also among the most enriched annotations. In contrast, potential crosstalk through Nodal and Hippo pathways showed very low enrichment in naive cells.

Having established transcriptional support for the TGF-β to SP1 cascade in naive cells, we asked how exhaustion reshapes it. We mapped differential expression between exhausted and naive CD8^+^ T cells onto the balanced cascades^80^ (Extended Data Fig. 4a,b, Methods and Table S5a,b). In the balanced cascade shared by the two shorter cores, 14 of 22 genes were significantly deregulated (Fig. 5e,f, Extended Data Fig. 4d, Methods and Table S5c). Receptors TGFBR2 and TGFBR3 decreased markedly, and SARA dropped strongly. SMADs shifted asymmetrically, with SMAD2 slightly increased, whereas SMAD4 and the negative feedback component SMAD7 decreased. Downstream of SMAD7, although the PP1 catalytic subunits (PPP1CA/CB/CC) increased, the PP1 adaptor GADD34 (PPP1R15A) decreased. Together, these changes indicated overall downregulation of genes encoding elements that support signal initiation at the receptor and downstream transduction, including significantly downregulated essential components such as TGFBR2 and SMAD4. In the longer alternatives, only four of the eleven additional genes were deregulated (Fig. 5f, Extended Data Fig. 4c,d, Methods and Table S5c). Notably, negative feedback components downstream of WWTR1 and SMAD7, such as SMURF1/2 and SKI, were downregulated. Given the low expression of the WWTR1-dependent segment in naive cells, the downregulation in exhaustion suggested that the additional segments of the longer alternatives were unlikely to carry signal.

BCEA corroborated these patterns, showing that the balanced cascade shared by the two shorter cores was more deregulated than that of the longer cores (Fig. 5g, Methods and Table S5d). Enrichment on the cores alone did not exceed that of their balanced counterparts, indicating deregulation within balancing species as well as the core. Thus, peripheral modules could contribute to deregulation of the TGF-β-SP1 axis beyond the core cascade. Across pathway annotations, deregulation concentrated in cell-cycle and DNA replication-repair programs, with additional shifts in metabolic responses and generic transcription (Fig. 5h, Extended Data Fig. 4e, Methods and Table S5e). This pattern linked the TGF-β-SP1 axis to altered proliferative control and genome maintenance in exhaustion. Deubiquitination was also highly deregulated, nominating disrupted SMAD4 recycling as a candidate balancing-derived mechanism that could alter transduction. In contrast, potential Nodal and Hippo crosstalk showed low deregulation, retaining the low expression observed in naive cells and reinforcing their limited engagement in exhaustion.

Overall, BCEA established that in naive CD8^+^ T cells, the TGF-β signaling cascade has transcriptional support for accessing metabolic programs via SP1 with intact enabling modules, whereas transcriptional changes in exhaustion limit this access. SP1 expression itself was not significantly altered in exhaustion (Extended Data Fig. 3f, 4f, Methods and Table S4f, S5f). This observation suggests a potential mechanism in which dampened upstream signaling reduces DNA binding and transcriptional output without loss of the TF itself, separating TF availability from TF effectiveness. Thus, BCEA results can guide downstream interpretation by linking cascade support and deregulation to specific TF regulatory interactions and, ultimately, to the metabolic pathways they control (Supplementary Fig. 9 and Table S4g, S5g).

## Discussion

This work presents SIGMA, a computational framework for analyzing signal transduction by converting curated prior knowledge into causally interpretable, elementally balanced cascades that connect defined sources to defined targets. Rather than enriching predefined pathway lists or fitting a single subnetwork to a particular dataset^6,8,12^, SIGMA enumerates feasible source-to-target cascades that respect information-flow logic and process-level constraints, and uses these cascades as the units for data interpretation. By enforcing elemental balance, SIGMA can connect processes across pathway catalogs in an unbiased manner, revealing links between modules not previously considered to interact. Enumerating mechanistic options matters biologically because cells rarely rely on a single canonical chain of reactions, and parallel routing contributes to robustness, crosstalk, and context specificity. By identifying the feasible set of cascades and interrogating them with data, SIGMA moves beyond traditional pathway enrichment to reveal which signaling cascades are most compatible with a given context and why.

Using SIGMA, we reconstructed a signaling network centered on TFs that regulate metabolism and built a receptor-TF roadmap. This map helps explain how conditioning cells with distinct ligands or microenvironments can drive similar metabolic outputs, and how the same cues can yield divergent outputs across contexts. It also nominates candidate nodes where targeted perturbations may rewire flow. We demonstrated the mechanistic resolution of SIGMA by connecting TGF-β to SP1, highlighting how elemental balance reveals supporting modules and potential crosstalk. Enumerating alternatives distinguished essential from cascade-specific species, pinpointing candidate bottlenecks and rerouting points. Process-level resolution of the cascades, together with essentiality, supports direct experimental translation by guiding reaction-level perturbations. Within SIGMA, we introduced BCEA to interpret measurements mechanistically, projecting omics data onto the extracted cascades and ranking alternatives. In CD8^+^ T cells^80^, BCEA revealed transcriptional support for the TGF-β-SP1 axis in naive cells that weakens with exhaustion, while localizing highly expressed or deregulated segments and quantifying crosstalk contributions. Mapping data to balanced cascades, rather than broad pathway labels, yields mechanistic, condition-specific hypotheses. Overall, this work lays the groundwork for integrating signaling and metabolic models within a unified framework, separating TF abundance from the upstream signaling capacity to regulate TF activity, and translating signals into regulatory constraints that may inform metabolic flux predictions^37,81–83^.

SIGMA can also be applied to single-cell analyses. Single-cell data reveal cell-state diversity and signaling rewiring across subpopulations, yet cluster-level summaries often conceal the mechanisms underlying these differences^84,85^. With SIGMA, data support for cascades could be compared across single cells and, by keeping alternatives explicit, cells shifting between viable cascades could be tracked. This analysis could enable a mechanistic understanding of subpopulation rewiring. Additionally, the framework can accommodate phosphoproteomics and other post-translational measurements as constraints on cascade activity^86,87^. Toward this application, reactions and species can be enabled, disabled, or prioritized based on post-translational modification evidence, thereby restricting the ensemble of cascades to those biochemically plausible in the condition of interest without refitting the network to the data.

In future work, elementally balanced cascades can serve as scaffolds for dynamical modeling^30–32,35,36^. Because each step specifies a biochemical process with defined stoichiometry, cascades can be translated directly into dynamical structures based on ordinary differential equations or Boolean logic, thereby reducing model space and limiting ad hoc wiring. In addition, SIGMA provides complete reaction sets and nominates kinetic and regulatory parameters to be estimated from time-resolved data or computational inference^88,89^.

Toward strengthening SIGMA, cascade quality will benefit from improvements in the granularity and completeness of the underlying knowledge base. Iterative curation can reduce cases where missing annotations omit valid cascades or overestimate essentiality^4,5,56^. BCEA also inherits biases from the data type used for projection, which can be addressed by broader integration of multi-omics datasets^90,91^, a future direction that SIGMA is designed to accommodate.

In summary, SIGMA reframes pathway analysis as elementally balanced cascade analysis. It enumerates alternative mechanistic cascades, ranks them against data, and exposes essential steps, crosstalk, and context-specific rewiring. In doing so, SIGMA converts prior knowledge and omics measurements into testable, process-level hypotheses. We anticipate applications across tissues, perturbations, single-cell datasets, and personalized contexts, where SIGMA may accelerate discovery and guide targeted interventions grounded in biochemical mechanisms.

## Methods

### Overview of SIGMA methodology

#### Signaling network representation

We represent the signaling network using binary incidence matrices derived from a PKN. Let m denote the number of directed reactions z_r_, with r ∈ {1, …, m}, and n the number of unique species x_s_, with s ∈ {1, …, n}. We define the reactant matrix R ∈ {0,1}^m×n^, where R_r,s_ = 1 if species x_s_ is a reactant in reaction z_r_. Similarly, the product matrix P ∈ {0,1}^m×n^ records the products of reactions, such that P_r,s_ = 1 if x_s_ is produced by z_r_. Regulatory inhibition is encoded in the matrix I ∈ {0,1}^m×n^, where I_r,s_ = 1 if species x_s_ inhibits reaction z_r_. To distinguish species that can transduce signal from cofactors and other small molecules, we define a Boolean vector c ∈ {0,1}^n^. An entry c_s_ = 1 indicates that species x_s_ is annotated, based on a curated list, as a cofactor or other small molecule that supports signal transduction but does not itself propagate the signal, whereas c_s_ = 0 indicates a signaling species. This representation establishes the precise role of each species in each reaction, providing a consistent basis for subsequent connectivity analysis and cascade extraction.

#### Core cascade extraction

We use two complementary approaches to extract core cascades that connect a specified source species to a target species: an algorithm based on graph search and an integer linear programming (ILP) formulation. Both approaches operate on a species-to-species adjacency matrix derived from the incidence matrices. To prevent extracting biologically irrelevant shortcuts through cofactors and small molecules, we first filter the network using the indicator vector c. For each species x_s_ with c_s_ = 1, we remove the corresponding column from R and P. This step yields a masked reactant matrix 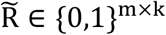 and a masked product matrix 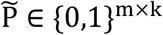, where k denotes the number of signaling species that are not annotated as cofactors or small molecules. We then compute the adjacency matrix A by matrix multiplication:

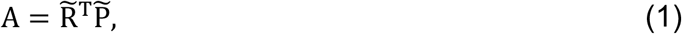

and subsequently convert it into a binary matrix A ∈ {0,1}^k×k^ by setting all entries greater than zero to one. An entry A_i,j_ = 1 indicates the existence of at least one reaction where species i is a reactant and species j is a product, while A_i,j_ = 0 indicates the absence of such a reaction. We exclude the inhibition matrix I from the adjacency construction to strictly model information flow rather than negative regulation.

##### Graph search

We convert the adjacency matrix A into an unweighted directed graph and apply breadth-first search to extract a species-to-species sequence with the shortest path length d_min_ between the specified source and target species. This directed species sequence defines a single signaling path of minimal length. To extract alternative paths of the same minimal length or longer, we enumerate all simple paths between the source and target with length between d_min_ and d_min_ + L_core_, where L_core_ ≥ 1 is a user-defined parameter that sets the maximum allowed number of additional steps beyond d_min_. We optionally restrict this enumeration to a predefined number of alternative paths.

##### Integer linear programming (ILP)

To extract directed signaling paths using optimization, we employ an ILP formulation adapted from the PathTracer algorithm developed by Tervo and Reed^92^. Connectivity is defined by binary decision variables 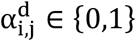, associated with the nonzero entries of the adjacency matrix A. For each pair of species with A_i,j_ = 1, the variable 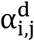 indicates whether the edge from species i to species j is active at step d of the path. We further define a termination variable S_d_ ∈ {0,1}, which indicates whether the path reaches the target species exactly at step d. The maximum search depth D is user-defined and is capped by the diameter of the network. We formulate the following ILP problem to ensure a valid path topology between a specified source-target pair:

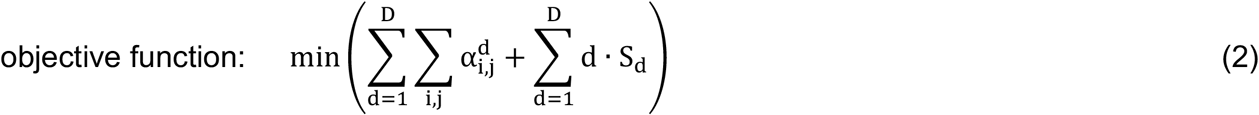

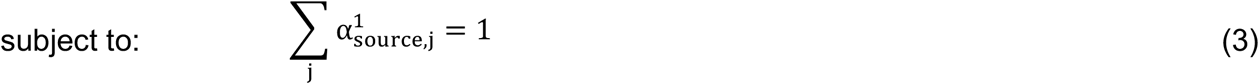

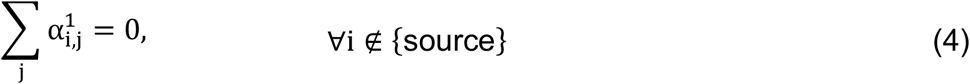

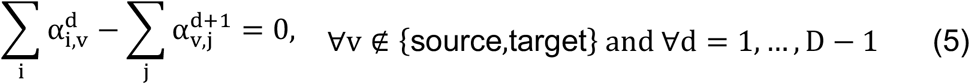

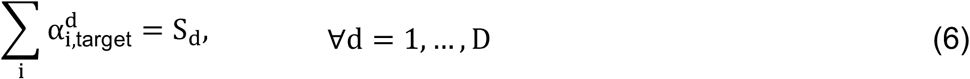

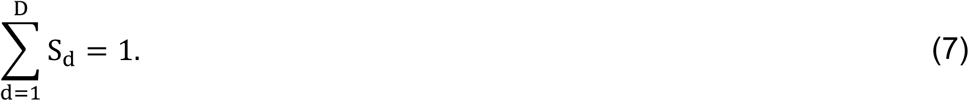

The objective function (2), subject to constraints (3)-(7), minimizes the sum of the number of active edges and the path length. Constraint (3) indicates that information flow must originate from the source at step d = 1. Consequently, constraint (4) ensures that no other species is allowed to transduce outgoing information flow at this initial step. For internal species v that are neither source nor target, we enforce information flow conservation between consecutive steps through constraint (5). Once the path reaches the target, information flow should not continue downstream. Accordingly, any information flow reaching the target at step d is captured by the termination variable in constraint (6). Finally, constraint (7) requires that the path reach the target exactly once within the maximum search depth D. Solving this ILP formulation yields a parsimonious information flow with minimal path length d_min_.

To extract alternative paths of the same minimal length or longer, we bound the maximum search depth to D^′^ = d_min_ + L_core_ and solve the ILP iteratively, up to a predefined number of alternatives. For each solution q, we define a set of indices for the active decision variables 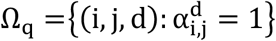. To ensure that subsequent solutions differ in topology from the previous solutions, we add an integer cut constraint that forces at least one decision variable that is active in solution q to become inactive in future solutions:

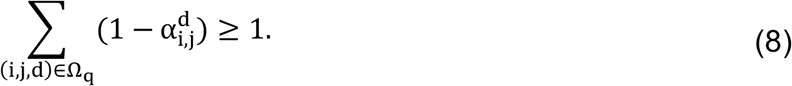

By accumulating such integer cuts over all previously found solutions, we enumerate a set of alternative species-to-species sequences that obey the same topological rules but differ in at least one connection at a specific step along the path, while still allowing the same connection to appear at a different step in another solution.

In contrast to the graph search procedure, the ILP formulation further supports guided navigation of the network. By adding linear constraints on selected edges, we can enforce or prohibit the activity of signaling components, incorporate known regulatory dependencies, and require consistency with activity states inferred from omics data. This flexibility enables us to test mechanistic hypotheses and observe how the set of feasible core paths changes.

##### Reaction-level core cascade reconstruction

Both graph search and ILP algorithms operate on the adjacency matrix A, which abstracts connectivity between species. To recover the biochemical reactions that can achieve this connectivity, we map each species-to-species sequence back to the incidence matrices 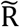 and 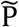. Let v_1_, v_2_, …, v_l_ denote the ordered sequence of species indices along a path from the source to the target. For each consecutive species transition, we identify the set of compatible reactions:

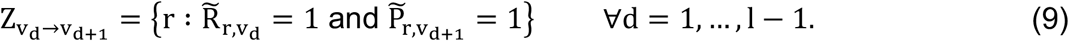

This set contains all directed reactions z_r_ that connect species 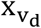 to species 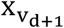. The ensemble of all concrete biochemical cascades that are consistent with the species sequence is then defined by the Cartesian product:

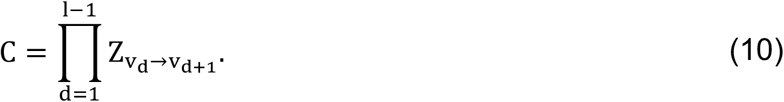

Each element of C represents a specific sequence of directed reactions that biochemically implements the corresponding species-to-species path. We apply this reconstruction procedure to every alternative path obtained from graph search or ILP, which yields the corresponding set of reaction-level core cascades for each source-target pair.

#### Balanced cascade extraction

To extract balanced signaling cascades from core cascades, we employ a second ILP formulation introduced by Mitsos et al.^26^ and further developed by Oftadeh and Marashi^93^, which we adapted to our SIGMA framework. This formulation translates the logical dependencies between reactions and species into integer constraints. It then optimizes the topology of active reactions subject to these constraints, while respecting causal information flow. In our framework, these logical dependencies arise directly from the reactant, inhibitor, and product incidence matrices R, I, and P derived from the PKN. The ILP formulation requires as input the reactions and species that are active in a given core cascade. We denote by R_core_ ⊆ {1, …, m} the set of indices of reactions that are active in the core cascade and by S_core_ ⊆ {1, …, n} the set of indices of species that are active in the core cascade. All reactions and species are associated with binary variables z ∈ {0,1}^m^ and x ∈ {0,1}^n^, where a value of one denotes activation and a value of zero denotes inactivation. We adapt the original ILP problem to our framework by formulating the following optimization problem:

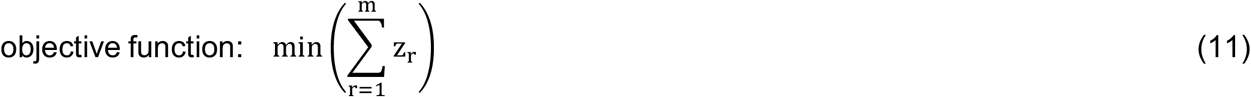

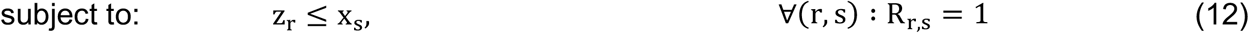

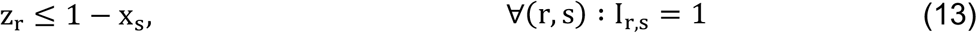

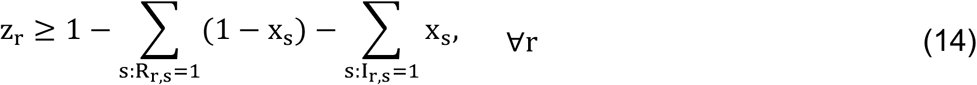

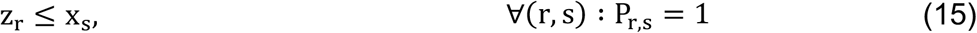

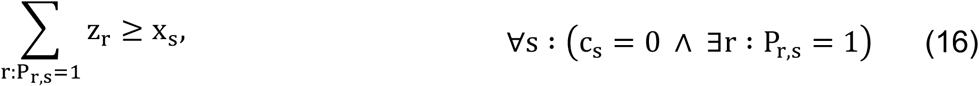

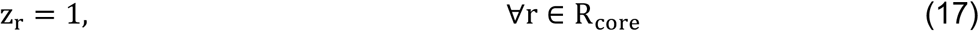

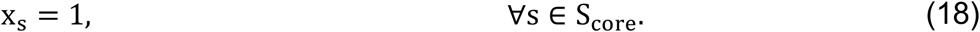

The objective function (11), subject to constraints (12)-(18), minimizes the number of active reactions needed to support the core cascade. Constraint (12) enforces that if a reaction is active, all its reactants must be active. Constraint (13) ensures that if a reaction is active, all its inhibitors must be inactive. When all reactants of a reaction are active and its inhibitors are inactive, the upstream conditions are sufficient for signal transduction, and constraint (14) enforces the reaction to be active so that the signal is propagated downstream in the network. Constraint (15) requires that if a reaction is active, all its products are active. For signaling species that are produced by at least one reaction, constraint (16) requires that their activity is supported by at least one active reaction producing them. We do not impose constraint (16) for species annotated as cofactors or other small molecules (c_s_ = 1), thereby preventing the introduction of artificial connections solely to balance the production of ubiquitous cofactors or small molecules that would not correspond to biologically meaningful cascades. Constraints (17) and (18) fix the reactions and species participating in the core cascade to be active. Solving this ILP formulation yields a minimal balanced cascade that supports signal transduction along the core cascade, with enabling reactions and participating species active and inhibitory regulators inactive.

After obtaining the minimal balanced cascade for a given core cascade by solving the ILP problem (11)-(18), we can further enumerate alternative balanced cascades. Let Z_min_ denote the number of active reactions in the minimal balanced cascade, as given by the objective value of the ILP. We then iteratively re-solve the ILP for balancing the same core cascade while enforcing that the objective value of any alternative solution does not exceed a user-defined tolerance above the minimum, that is, Z_min_ + L_bal_, where L_bal_ is the size relaxation parameter that controls the maximum allowed increase in the number of active reactions relative to the minimal balanced cascade. To ensure that each additional solution corresponds to a distinct balanced cascade, we introduce integer cut constraints that exclude previously found reaction activity patterns. For each solution p with reaction activity vector z^(p)^, we define the index set of active reactions 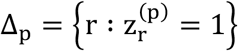. For all subsequent solutions, we impose the constraint:

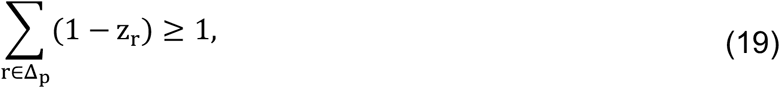

which forces at least one reaction that is active in solution p to become inactive in any new solution. By repeatedly solving the ILP with the accumulated integer cuts (19), up to a predefined maximum number of alternatives, we obtain a family of distinct balanced cascades that support signal transduction along the same core cascade while differing by at least one active reaction.

#### Balanced cascade enrichment analysis

To assess whether balanced cascades are enriched for expressed or deregulated species, we perform BCEA using transcriptomic or proteomic measurements. BCEA can use absolute or differential omics data from bulk or single-cell experiments, depending on the analysis. We employ an over-representation analysis based on the upper tail of the hypergeometric cumulative distribution to evaluate whether a cascade contains more expressed or deregulated species than expected by chance. Because inhibitors in a balanced cascade are constrained to be inactive, we exclude inhibitory species from the cascade sets used in the enrichment analysis. However, we can assess their expression or deregulation separately to enable inspection of potential cascade inhibition.

##### Absolute omics

For each experimental condition, we require an omics dataset indexed by gene or protein identifiers (for example, UniProt identifiers). For absolute gene expression data, we use normalized values that enable comparisons across genes within a condition, for example, TPM values for bulk data and counts per 10,000 (CP10k) for single-cell data. Similarly, for proteomics, we use quantitative protein abundance measures that are comparable across proteins within a condition. We define O^abs^ as the set of gene or protein identifiers (IDs) measured in the absolute dataset. The subset of expressed genes or proteins is E^abs^ = {ID ∈ O^abs^ : expression(ID) ≥ θ_expr_}, where θ_expr_ is a user-defined expression threshold, above which a gene or protein is considered active. For enrichment, we restrict the background to genes or proteins present in both the PKN and the omics dataset. Let U_PKN_ denote the set of all identifiers that map to PKN species and define the background as B^abs^ = O^abs^ ∩ U_PKN_. The expressed background is 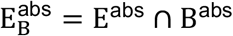. For each balanced cascade c with associated set of gene or protein identifiers S_c_, we retain only those identifiers that are present in the background, 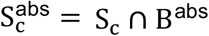, and define the set of expressed identifiers in the cascade as 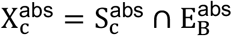. We compute the enrichment p-value for the balanced cascade c from the upper tail of the hypergeometric cumulative distribution as:

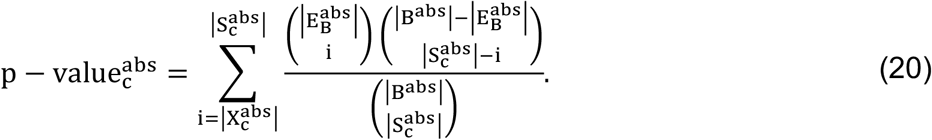

Here ∣ B^abs^ ∣ is the size of the background, 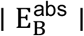 is the number of expressed background identifiers, 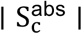 is the number of cascade identifiers found in the background, and 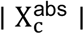 is the number of expressed background identifiers in the cascade. We quantify the fraction of expressed genes or proteins in the balanced cascade c as:

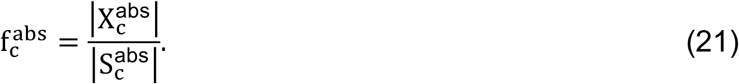

##### Differential omics

In the differential setting, both transcriptomics and proteomics datasets are typically provided as fold changes, together with adjusted *P* values, for the compared conditions. We define O^diff^ as the set of gene or protein identifiers measured in the differential dataset. The subset of significantly deregulated genes or proteins is E^diff^ = {ID ∈ O^diff^ : (FC(ID) ≥ θ_FC_ ∨ FC(ID) ≤ 1/θ_FC_) ∧ adjP(ID) ≤ θ_adjP_}, where θ_FC_ is a user-defined fold change threshold and θ_adjP_ is a user-defined adjusted *P* value cutoff. As in the absolute omics case, we restrict the background to genes or proteins that appear both in the PKN and in the differential dataset B^diff^ = O^diff^ ∩ U_PKN_, and define the deregulated background as 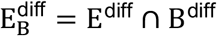. For each balanced cascade c with associated set of identifiers S_c_, we retain only identifiers that belong to the background 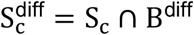. We define the set of deregulated identifiers in the cascade as 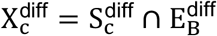. We calculate a p-value that quantifies whether the balanced cascade c is enriched for deregulated identifiers using the upper tail of the hypergeometric cumulative distribution as:

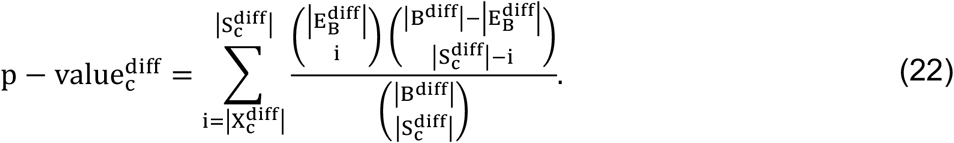

Here ∣ B^diff^ ∣ is the size of the background, 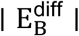 is the number of deregulated background identifiers, 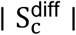 is the number of cascade identifiers found in the background, and 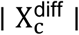 is the number of deregulated background identifiers in the cascade. We quantify the fraction of deregulated genes or proteins in the balanced cascade c as:

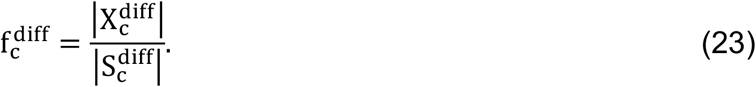

### Reactome signaling network assembly

We downloaded *Homo sapiens* pathways in Systems Biology Markup Language (SBML) format from Reactome (version 90)^56^. We parsed the SBML files in Python (version 3.9.6) to extract the species and reactions associated with each Reactome pathway. For each species, we retained the Reactome identifier, species name, entity type, pathway membership, and available UniProt, Ensembl and ChEBI identifiers. Species with protein components were mapped to one or more UniProt identifiers. These mappings allowed Reactome species, including complexes, to be linked to the genes encoding their protein components. For each reaction, we retained the Reactome identifier, reaction name, pathway membership, and Enzyme Commission (EC) numbers.

We organized the parsed Reactome information by representing reactions and species as sparse incidence matrices, following the signaling network representation described above. The reactant matrix included species annotated in Reactome as reactants, activators, or catalysts, because these species represent upstream requirements for reaction activity. Product and inhibitor matrices were stored separately. This Reactome signaling network contained 15,180 reactions, 25,422 species, 11,616 genes, and 2,742 pathways, and served as the starting point for reconstruction of the TF-specific signaling network.

### TF-specific signaling network reconstruction

#### TFs regulating metabolism

To identify TFs regulating metabolism, we employed the human genome-scale model Recon3D^54^, which contains 5,835 metabolites and 10,600 reactions. These reactions are associated with 2,248 metabolic genes, including isoform entries, that encode metabolic enzymes. From this list of metabolic genes, we extracted 1,884 unique entries and mapped them to TRRUST (version 1)^55^, a literature-curated resource of human transcriptional regulatory interactions that contains 8,015 regulatory relationships for 748 TFs. TRRUST annotates not only classical DNA-binding regulators as transcription factors but also transcriptional co-regulators and components of the general transcription machinery. We retained only regulatory relationships annotated as activation or repression and excluded relationships with unknown mode of action. Mapping resulted in 460 regulatory relationships between 115 TFs and 237 metabolic genes (Table S1a). Using the gene-protein-reaction (GPR) rules stored in Recon3D, we also mapped each of the 237 metabolic gene targets to the metabolic reactions catalyzed by the encoded enzyme or enzymatic complex. Because each reaction in Recon3D is assigned to a metabolic subsystem, this mapping linked each TF to the metabolic pathways regulated by its target genes.

#### TF states in Reactome

We mapped the TFs that regulate metabolism to species in the assembled Reactome network. If a species contained at least one of the UniProt identifiers of the 115 TFs and was localized to the nucleus but not bound to a specific DNA segment, we classified it as a TF state. TF states can include complexes of multiple TFs, so the same state can be assigned to different TFs. In total, 101 TFs had corresponding entries in Reactome, yielding 532 unique TF states.

#### Pathway expansion around metabolism

We used these 532 TF states as root species and mapped them to Reactome pathways. Pathways containing at least one root species were defined as the first layer. Using the visual links in the Reactome online pathway browser (https://reactome.org/PathwayBrowser/), which denote upstream or downstream relationships between pathways (inter-pathway connectivity), we identified all pathways connected to the first layer. Newly identified pathways that were not already included in the first layer were defined as the second layer. We repeated this expansion for two additional steps, resulting in four expansion layers and 268 unique pathways (Table S1d).

#### Gap filling and network assembly

We performed gap filling across the 268 collected pathways. For each pathway, we annotated intra-pathway reactions by visually inspecting dashed links in the Reactome online pathway browser (https://reactome.org/PathwayBrowser/). These links connect an entity to a group of functionally equivalent proteins or small molecules that either includes the entity or overlaps with it. Because these links lack a reported directionality, we included both directions for each one. In total, we added 1,439 unique directional reactions for intra-pathway links between single species. We also traced directed pathway-pathway connections in Reactome, shown as green-and-white boxes in the online pathway browser, and added 69 inter-pathway reactions when the linked pathways had no species overlap. For each such link, we formed a reaction by combining one or more species from the source pathway as reactants and one or more species from the target pathway as products, and we consulted the literature when the linkage was not obvious. During gap filling, we also introduced 193 translocation events to link compartmentalized forms of the same species when the same species name appeared in distinct Reactome pathways. We represented each event as two split directional reactions, generating a total of 386 reactions. After gap filling, we merged the 268 curated pathways into the incidence-matrix representation described above, yielding the final TF-specific signaling network comprising 9,227 reactions, 12,288 species, and 6,372 genes.

#### Receptor identification

We identified plasma membrane receptors and their ligands in an unbiased manner. We collected all reactions that contained at least one extracellular species and at least one plasma membrane species as reactants. To avoid collecting transport reactions instead of ligand-receptor binding reactions, we checked whether the products of the reaction included at least one of the plasma membrane proteins, implying the formation of a ligand-receptor complex as a product of the binding reaction. Using this reaction-collection procedure, we extracted 224 receptors, 227 ligands, and 319 ligand-receptor interactions in the TF-specific signaling network (Table S1e).

#### Cofactor and small molecule identification

For every species in the assembled Reactome network, we computed degree centrality, which denotes the number of other species to which it is connected^59^. Among the nodes with a degree centrality greater than 20 (top 1%), we manually identified species corresponding to cofactors and small molecules. We ensured complete coverage of nucleotide cofactors (NTPs, NDPs, and NMPs) by adding those not captured by this high-degree set. The final curated list comprised 54 cofactor and small molecule entities (Table S1h), mapped to one or more compartmentalized species in the Reactome and TF-specific signaling networks. Classical second messengers that operate as signaling molecules, namely cAMP, cGMP, IP3, DAG, and Ca^2+^, were not included in this list and were retained in the core cascade extraction^94^.

#### Effect of cofactors and small molecules on network topology

To assess the topological impact of cofactors and small molecules, we compared the TF-specific signaling network before and after removing the 54 curated entities. For each network, we computed the degree centrality of every species and used the resulting degree distribution to quantify the effect of removing cofactors and small molecules on local connectivity. The average degree centrality was 4.1 before removal and 3.1 after removal. We also computed shortest-path distances between all reachable pairs of species in the directed species graphs. From these distances, we calculated the frequency of each path length and defined the network diameter as the longest shortest-path distance among reachable species pairs. The network diameter was 40 before removal and 105 after removal.

### Receptor-to-TF connectivity mapping

#### Receptor-to-TF roadmap

To construct the connectivity roadmap between receptors and TFs in the TF-specific signaling network, we used the 224 plasma membrane receptors as source species and the 532 TF states as target species. For each receptor-TF state pair, we extracted the shortest core cascade using the graph search algorithm described above. A receptor was considered linked to a TF if a directed path connected the receptor to at least one Reactome state of that TF. Because a single TF can correspond to multiple Reactome states, we summarized path lengths across the linked receptor-TF state pairs as a single value for each receptor-TF pair. Thus, when we report the shortest path length between a receptor and a TF, this value refers to the average shortest path length from that receptor to all reachable Reactome states assigned to the TF (Extended Data Fig. 2a and Table S2a).

#### Receptor and TF families

We assigned receptors and TFs to higher-level families using the pathway hierarchy relationships downloaded from Reactome (version 90)^56^. For each of the 224 receptors, family assignment was based on the pathways in the TF-specific network that contained the receptor species. For each of the 101 TFs, assignment was based on the pathways containing the TF states assigned to that TF. Because a receptor or TF could appear in more than one Reactome family, the same receptor or TF could contribute to multiple family counts. For example, a receptor or TF appearing in two families contributed one count to each family. For each receptor family, we counted both the total number of receptors assigned to the family and the number of those receptors linked to at least one TF that regulates metabolic genes. Similarly, for each TF family, we counted both the number of TFs regulating metabolic genes assigned to the family and the number of those TFs linked to at least one receptor. To compare receptor families by their proximity to TFs, we built a path-length profile for each receptor family, containing one distance value between that receptor family and each linked TF. For a given receptor family and TF, we defined this distance as the average receptor-TF shortest path length across the receptors in that family that were linked to that TF. The receptor-TF shortest path lengths used for this average were defined in the receptor to TF roadmap above. We also used these path-length profiles to cluster the receptor families using Euclidean distance (Extended Data Fig. 2b and Supplementary Fig. 1b).

#### Receptor and TF subnetwork reconstruction

For each connected receptor-TF state pair, we exhaustively enumerated all core cascades with length equal to the shortest path length for that pair using the graph search algorithm. For each alternative core cascade, we extracted all corresponding minimal-size balanced cascade alternatives using the ILP formulation described above. Using the enumerated core and balanced alternatives, we represented each receptor-TF state pair as a species-frequency profile. For each species, its frequency was defined as the fraction of alternative cascades for that receptor-TF state pair in which the species appeared. A frequency of one indicated that the species appeared in all alternatives for that pair, whereas a frequency of zero indicated that it appeared in none. For each receptor-TF state pair, we calculated one frequency profile using only the core alternative cascades and one using the balanced alternative cascades. For balanced cascades, we included core species and supporting species added during balancing, whereas inhibitory species constrained to be inactive were not included in these frequency profiles. These profiles served as the basis for reconstructing receptor and TF subnetworks.

For each TF linked to at least one receptor, we reconstructed its upstream subnetwork as the union of cascades from connected receptors to the reachable TF states assigned to that TF. Each species in the subnetwork was weighted by its average frequency across all species-frequency profiles corresponding to connected receptor-TF state pairs assigned to that TF. If one TF state was linked to multiple receptors, each link contributed as a separate frequency profile to the average. Similarly, if one receptor was connected to multiple states of the same TF, each connection also contributed separately. We performed this subnetwork reconstruction once for the core and once for the balanced species-frequency profiles, yielding one core upstream subnetwork and one balanced upstream subnetwork for each TF.

For each receptor linked to at least one TF, we reconstructed its downstream subnetwork as the union of minimal balanced cascades from that receptor to its reachable TF states. Each species in the subnetwork was weighted by its average frequency across all species-frequency profiles corresponding to connected receptor-TF state pairs involving that receptor. If the receptor was connected to multiple TF states, including multiple states assigned to the same TF, each connection contributed separately to the average. For this reconstruction step, we used only the balanced species-frequency profiles.

#### Global species essentiality and subnetwork similarity

The global species essentiality score quantified how broadly and consistently a species contributed to the receptor-mediated reachability of TFs regulating metabolic genes. To calculate global species essentiality, we used the frequency-weighted TF-upstream subnetworks defined in the preceding section. For each species, we averaged its subnetwork weight across all TF-upstream subnetworks. When a species was absent from a TF-upstream subnetwork, its weight for that subnetwork was set to zero before averaging. We calculated one global species essentiality score using the core TF-upstream subnetworks and one using the balanced TF-upstream subnetworks (Fig. 3c and Table S2e).

To quantify subnetwork similarity, we performed pairwise comparisons among balanced receptor-downstream subnetworks and among balanced TF-upstream subnetworks defined in the preceding section. Each subnetwork was represented as a vector containing the weights of its species. Vectors were defined over a common species set detected across the balanced subnetworks, with species absent from a given subnetwork assigned a weight of zero in that vector. We computed one receptor-receptor similarity matrix from pairwise Spearman rank correlations between receptor-downstream subnetwork vectors and one TF-TF similarity matrix from pairwise Spearman rank correlations between TF-upstream subnetwork vectors. We used the same species-weight vectors to cluster balanced receptor-downstream and balanced TF-upstream subnetworks using Spearman distance (Fig. 3d,e and Supplementary Fig. 3).

### TGF-β to SP1 cascade reconstruction

#### Cascade extraction

To extract the balanced signaling cascades connecting the TGF-β receptor to SP1, we used TGFBR2 in the plasma membrane as the source species and p-2S-SMAD2,3:SMAD4:SP1 in the nucleoplasm as the target species. Although SP1 has four states in Reactome, p-2S-SMAD2,3:SMAD4:SP1 was the only SP1 state connected to any of the identified receptors. We extracted the shortest core cascade and exhaustively enumerated alternative core cascades by setting L_core_ = 105, corresponding to the diameter of the TF-specific network after cofactor and small molecule removal. This enumeration identified four alternative core cascades. For each alternative core cascade, we extracted the corresponding minimal-size balanced cascade using the ILP formulation. We set L_bal_ = 0 to enumerate additional balanced cascades with the same minimal size obtained from balancing each core cascade, but did not identify any additional feasible alternatives for any of the four core cascades.

#### Cascade analysis

To identify essential species and reactions that sustain TGF-β signaling to SP1, we computed the intersections of species sets and reaction sets across the four balanced cascade alternatives. Species or reactions present in all four balanced cascades were considered essential for signal transduction between this source and target, whereas those present in only a subset of alternatives were considered alternative-specific.

We mapped reactions in each alternative balanced TGF-β to SP1 cascade to Reactome pathway annotations. For each alternative cascade, we calculated the fraction of cascade reactions annotated to each pathway represented in that cascade. If a reaction was annotated to more than one pathway, it contributed once to each pathway. For each represented pathway, we further partitioned the set of cascade reactions mapped to that pathway into core reactions and reactions added during balancing, and calculated their relative fractions within that set. To quantify the coverage of each represented Reactome pathway by each alternative cascade, we divided the number of cascade reactions mapped to that pathway by the total number of reactions annotated to that pathway in Reactome.

To estimate the energetic demand of each alternative balanced TGF-β to SP1 cascade, we identified ATP-consuming reactions and used the ATP stoichiometry annotated in Reactome for each reaction. We computed the total ATP requirement of each cascade by summing the stoichiometric ATP consumption across the identified reactions in that cascade. All four balanced cascade alternatives contained the same ATP-consuming reactions and therefore had identical total ATP requirements.

### Mapping transcriptomic data onto TGF-β to SP1 cascades

#### Naive CD8^+^ T cell expression

Three independent naive CD8^+^ T cell samples (Table S4a), publicly available in Gene Expression Omnibus (GEO) under accession number GSE227316^80^, were used to quantify absolute expression for the analysis of TGF-β to SP1 signaling. We converted the available raw count matrix to transcripts per million (TPM) using gene lengths obtained from BioMart (version 113). Only genes present in both the count matrix and the gene-length annotation list were retained. For each sample, raw counts were divided by gene length in kilobases, yielding reads per kilobase (RPK). RPK values were then normalized by the sample-specific sum of RPK values and multiplied by 10^6^ to derive TPM values. We averaged TPM values across the three naive samples to obtain a single representative absolute expression value for each Ensembl gene identifier. For subsequent mapping of gene expression values to Reactome species, we converted Ensembl identifiers to UniProt identifiers using an Ensembl-to-UniProt mapping downloaded from BioMart (version 113). We retained only genes present in both the averaged TPM table and the BioMart mapping table and excluded matched entries without a UniProt identifier. When one Ensembl identifier mapped to multiple UniProt identifiers, the same TPM value was assigned to each UniProt identifier. When multiple Ensembl identifiers mapped to the same UniProt identifier, we averaged their TPM values into one expression value per UniProt identifier. These UniProt-level TPM values (Table S4b) were used to map absolute expression onto the alternative cascades and to perform BCEA.

#### Absolute expression mapping and enrichment across TGF-β to SP1 cascades

Based on the distribution of TPM values in naive CD8^+^ T cells (Extended Data Fig. 3b), we classified genes into four categories: highly expressed genes (TPM ≥ 10), moderately expressed genes (1 ≤ TPM < 10), genes with low expression (0 < TPM < 1), and genes with no detected expression (TPM = 0). For each alternative TGF-β to SP1 cascade, we performed BCEA at the gene level by collecting all genes annotated to the Reactome species participating in that cascade (Table S3c,d). We used θ_expr_ = 10 as the expression threshold for considering genes active, thereby selecting genes in the highly expressed category (TPM ≥ 10). We calculated the absolute enrichment p-value, 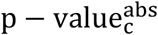, and expressed fraction, 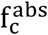, for each of the four alternative core cascades and each of the four corresponding balanced cascades. We adjusted these eight absolute enrichment p-values together across the core and balanced alternatives using Benjamini-Hochberg correction.

#### Reactome pathway enrichment and TF analysis with absolute data

We performed pathway enrichment analysis at the gene level for each of the 268 Reactome pathways that constituted the TF-specific network, using the genes annotated to each pathway. To calculate enrichment from the naive CD8^+^ T cell absolute expression data, we used over-representation analysis with the upper tail of the hypergeometric cumulative distribution and adjusted the resulting p-values across all 268 pathways using Benjamini-Hochberg correction. Genes with TPM ≥ 10 defined the expressed gene set for this enrichment analysis. To assess TF expression in naive CD8^+^ T cells, we mapped the UniProt identifiers of the 115 TFs that regulate metabolic genes to their TPM values.

#### Exhausted versus naive CD8^+^ T cell differential expression

To perform differential gene expression analysis between exhausted and naive CD8^+^ T cells, we used the three naive CD8^+^ T cell samples described above and three exhausted CD8^+^ T cell samples from the same study^80^ (Table S5a). The exhausted cells from that study were repeatedly antigen-stimulated T cells that phenocopied chronic antigen stimulation and exhaustion. Both naive and exhausted sample groups are publicly available in GEO under accession number GSE227316^80^. We used the available raw count matrices for the naive and exhausted samples to perform differential gene expression analysis with the DESeq2 R package (version 1.38.3). This analysis yielded log_2_ fold changes and Benjamini-Hochberg adjusted *P* values for exhausted versus naive CD8^+^ T cells. For subsequent mapping of differential gene expression values to Reactome species, we converted Ensembl identifiers to UniProt identifiers using the Ensembl-to-UniProt mapping downloaded from BioMart (version 113). We retained only genes present in both the DESeq2 output and the BioMart mapping table and excluded matched entries without a UniProt identifier. When one Ensembl identifier mapped to multiple UniProt identifiers, the same differential expression values were assigned to each UniProt identifier. When multiple Ensembl identifiers mapped to the same UniProt identifier, we averaged their log_2_ fold changes and combined their adjusted *P* values using the harmonic mean. These UniProt-level log_2_ fold changes and adjusted *P* values (Table S5b) were used to map differential expression onto the alternative TGF-β to SP1 cascades and to perform differential BCEA.

#### Differential expression mapping and enrichment across TGF-β to SP1 cascades

For differential BCEA at the gene level, each alternative TGF-β to SP1 cascade was represented by all genes annotated to its participating Reactome species (Table S3c,d). Using the exhausted versus naive CD8^+^ T cell differential expression data, we performed BCEA with θ_FC_ = 1.3 as the fold change threshold and θ_adjP_ = 0.05 as the adjusted *P* value cutoff to define significantly deregulated genes. We calculated the differential enrichment p-value, 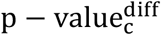, and deregulated fraction, 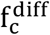, for each of the four alternative core cascades and each of the four corresponding balanced cascades. We adjusted these eight differential enrichment p-values together across the core and balanced alternatives using Benjamini-Hochberg correction.

#### Reactome pathway enrichment and TF analysis with differential data

For differential Reactome pathway enrichment analysis at the gene level, each of the 268 pathways constituting the TF-specific network was represented by its annotated genes. To calculate enrichment from the exhausted versus naive CD8^+^ T cell differential expression data, we used over-representation analysis with the upper tail of the hypergeometric cumulative distribution and adjusted the resulting p-values across all 268 pathways using Benjamini-Hochberg correction. Genes with |log_2_(fold change)| ≥ log_2_(1.3) and adjusted *P* value ≤ 0.05 defined the deregulated gene set for this enrichment analysis. To assess TF differential expression between exhausted and naive CD8^+^ T cells, we mapped the UniProt identifiers of the 115 TFs that regulate metabolic genes to their log_2_ fold changes and adjusted *P* values.

### Software and visualization

Data plotting and curve fitting were primarily performed in GraphPad Prism (version 11.0.0). Hierarchical clustering and dendrogram plotting were performed in R (version 4.2.2). Sankey diagrams and row z-scored expression heatmaps were also generated in R. Large-scale networks were visualized with Cytoscape (version 3.10.2). Enrichment tests and Benjamini-Hochberg corrections were performed in MATLAB (version R2024b). Figures and layouts were refined in Adobe Illustrator 2022 (version 26.3.1), which was also used to create the schematic illustrations and alternative TGF-β to SP1 cascade diagrams included in the manuscript.

## Code and data availability

The SIGMA code, data, and results supporting this study are publicly available at https://github.com/EPFL-LCSB/sigma. The repository enables users to: 1) reproduce the analyses and outputs reported in the manuscript; 2) use the provided Reactome network or curated TF-specific network to connect species of interest; and 3) execute the SIGMA workflow from scratch to generate a new network centered on user-defined root species and perform additional case studies. SIGMA was developed and tested with MATLAB R2024b and uses IBM ILOG CPLEX (version 12.10) as the optimization solver.

## Acknowledgements

This work was supported by funding from the Swiss National Science Foundation grants 188623, 198543 (Sinergia), 180575 and 225148 (NCCR), and by the École Polytechnique Fédérale de Lausanne (EPFL). ChatGPT-5 was used to polish grammar and syntax in this manuscript. The authors reviewed and edited the text and take responsibility for the final manuscript.

## Contributions

Conceptualization: D.L., M.M., and V.H.; methodology: D.L., O.O., M.T., M.M., and V.H.; software: D.L., O.O., M.T., and M.M.; data curation: D.L., O.O., M.T., and M.M.; formal analysis: D.L.; visualization: D.L.; writing - original draft: D.L.; writing - review and editing: D.L., M.M., and V.H.; supervision: M.M. and V.H.; funding acquisition: V.H.

## Competing interests

The authors declare no competing interests.

## Extended Data Figures

**Extended Data Fig. 1.**
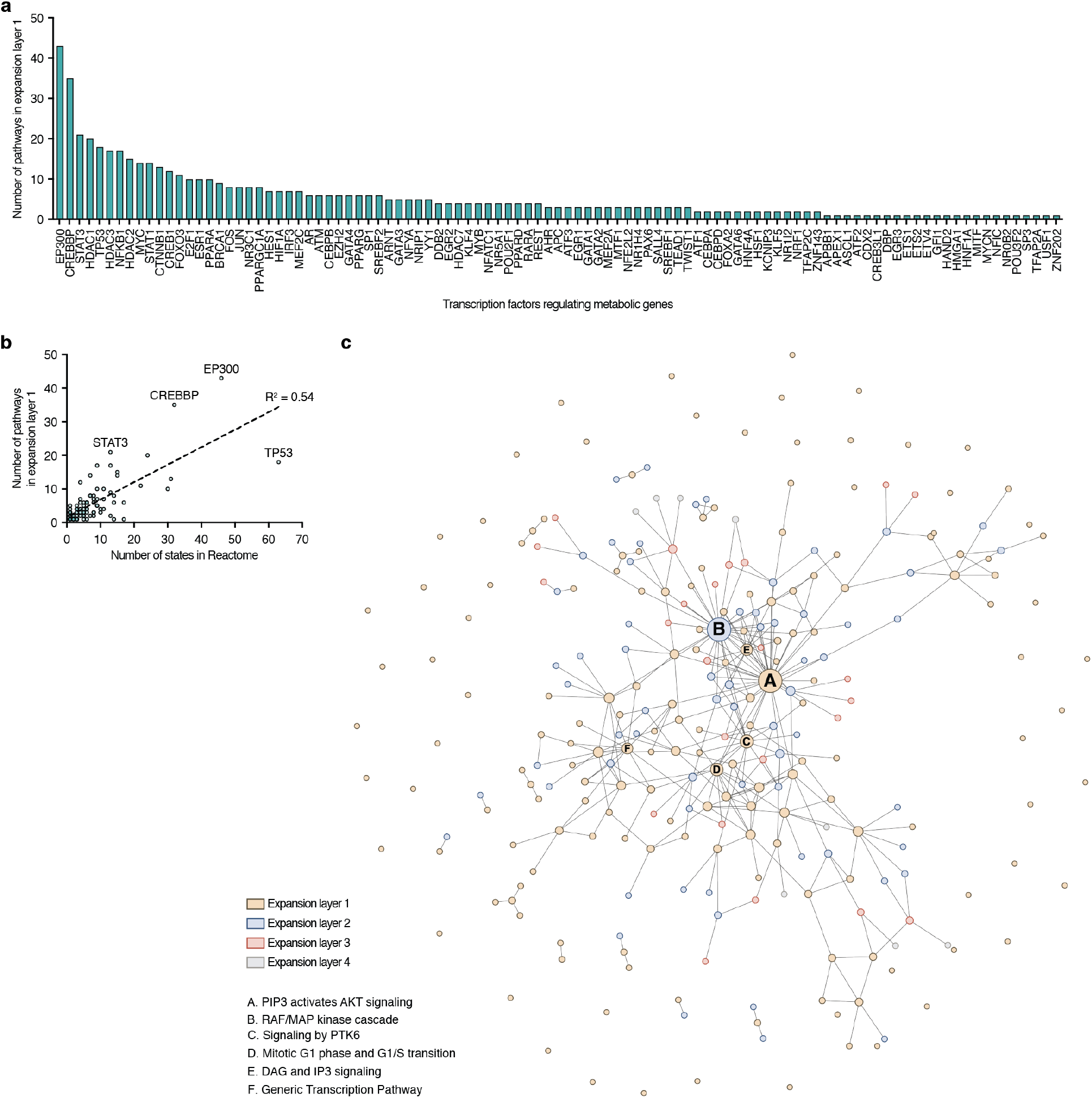
Reactome pathway coverage and connectivity for transcription factors regulating metabolism. **a**, Bar plot showing transcription factors (TFs; n = 101) that regulate metabolic genes and are mapped to at least one Reactome pathway. Bars are ordered by the number of Reactome pathways containing each TF at expansion layer 1. **b**, Correlation between the number of Reactome states per TF and the number of expansion layer 1 pathways containing that TF (n = 101). The dashed line shows a simple linear regression fit (R^2^ = 0.54). **c**, Network view of the pathway collection expanded across four layers from TF root species. Nodes represent Reactome pathways (n = 268) and edges denote documented inter-pathway links. Highly interconnected pathways are labeled (lettered IDs). Figure created with Cytoscape.

**Extended Data Fig. 2.**
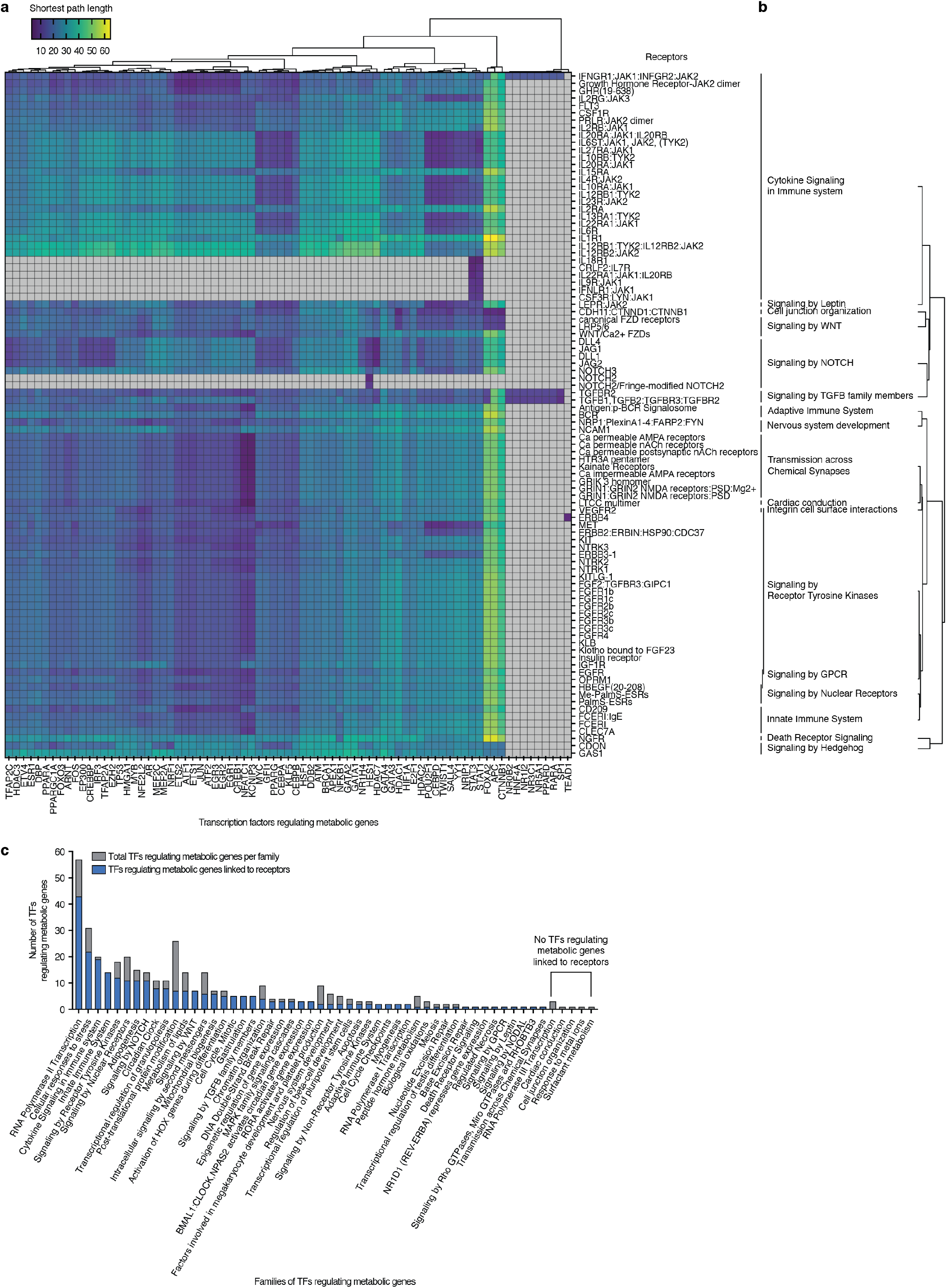
Receptor-to-transcription factor connectivity for metabolic regulation. **a**, Heatmap showing the shortest-path lengths connecting receptors (n = 93) to TFs that regulate metabolic genes (n = 77). Gray boxes indicate no connection for that receptor-TF pair. Hierarchical clustering of TFs is also shown (Euclidean distance, Ward’s linkage). **b**, Mapping of receptors linked to TFs that regulate metabolic genes (n = 93) to their corresponding receptor families, with hierarchical clustering of the families (Euclidean distance, Ward’s linkage). **c**, Number of TFs regulating metabolic genes per TF family in the TF-specific network (gray) and number linked to receptors (blue). Bars are ordered by the count of linked TFs regulating metabolic genes. Five TF families had no linked TFs regulating metabolic genes (right).

**Extended Data Fig. 3.**
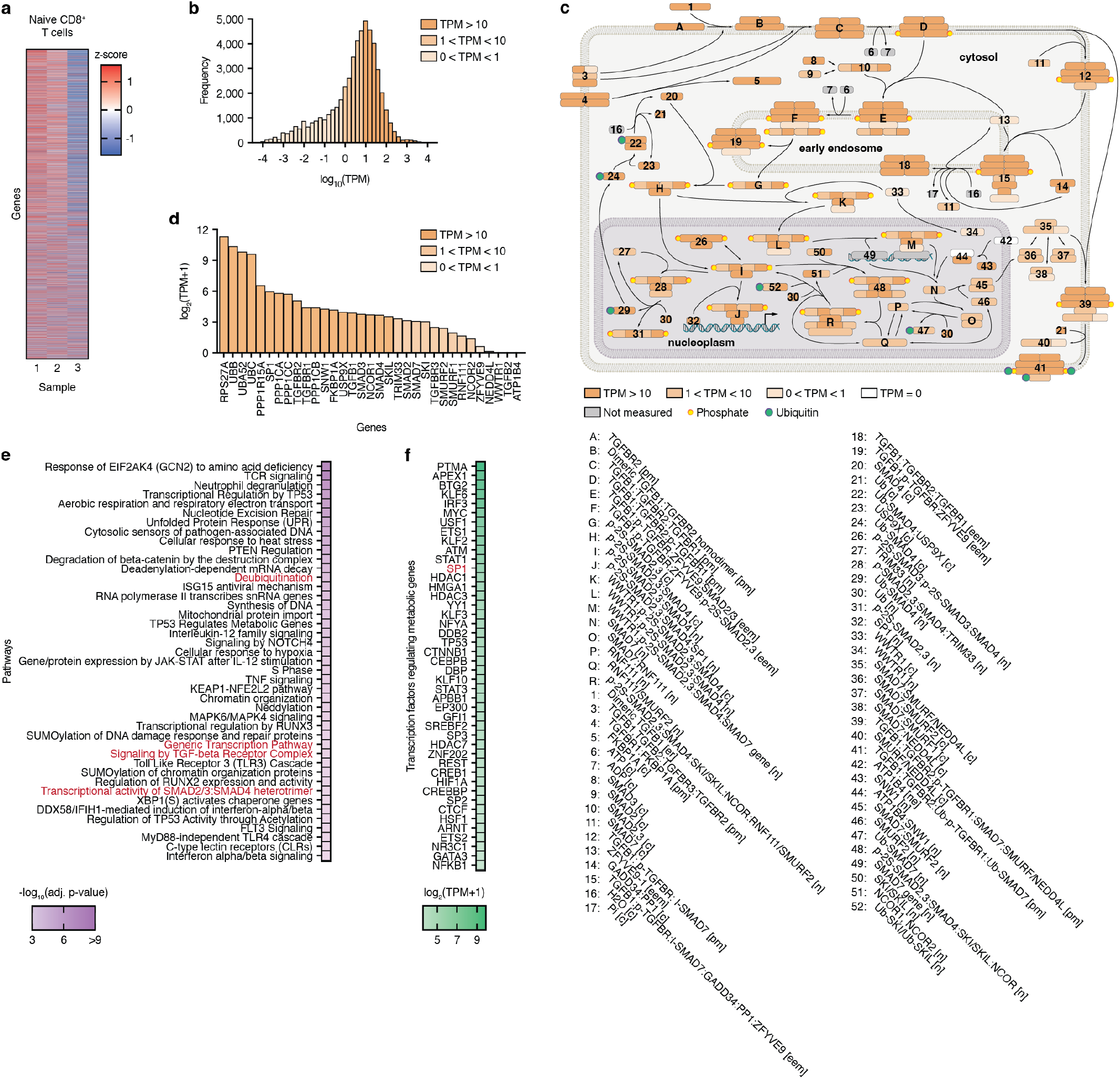
Naive CD8^+^ T cell expression and mapping onto TGF-β to SP1 cascades. **a**, Heatmap of normalized expression, z-scored by row, for naive CD8^+^ T cell samples (n = 3) used in this study, publicly available in Gene Expression Omnibus (GEO) under accession number GSE227316. **b**, Histogram of absolute expression levels (TPM: transcripts per million) for the naive CD8^+^ T cell dataset. The x-axis shows log_10_(TPM). Bar color intensity (orange) indicates TPM ranges. **c**, Mapping of absolute expression from naive CD8^+^ T cells onto the balanced cascade shared by the 17- and 20-step cores. Color intensity (orange) corresponds to TPM for each UniProt-mapped gene represented in the cascade species (Methods). Protein-complex species show separate color intensities for the UniProt-mapped gene associated with each constituent protein. White indicates undetected genes. **d**, Bar plot showing gene expression in naive CD8^+^ T cells for all genes appearing across the TGF-β to SP1 alternative cascades. The y-axis shows log_2_(TPM + 1). Bar color intensity (orange) indicates TPM ranges as in panels **b** and **c. e**, Heatmap showing the most enriched Reactome pathway annotations participating in the TF-specific network (adjusted p-value ≤ 0.001), calculated from absolute expression. Enrichment was performed at the gene level with over-representation analysis using the upper tail of the hypergeometric cumulative distribution and Benjamini-Hochberg correction. Pathways present in the TGF-β to SP1 alternative cascades are labeled in red. **f**, Heatmap showing the most expressed TFs that regulate metabolic genes (TPM ≥ 10), calculated from absolute expression. SP1 is labeled in red.

**Extended Data Fig. 4.**
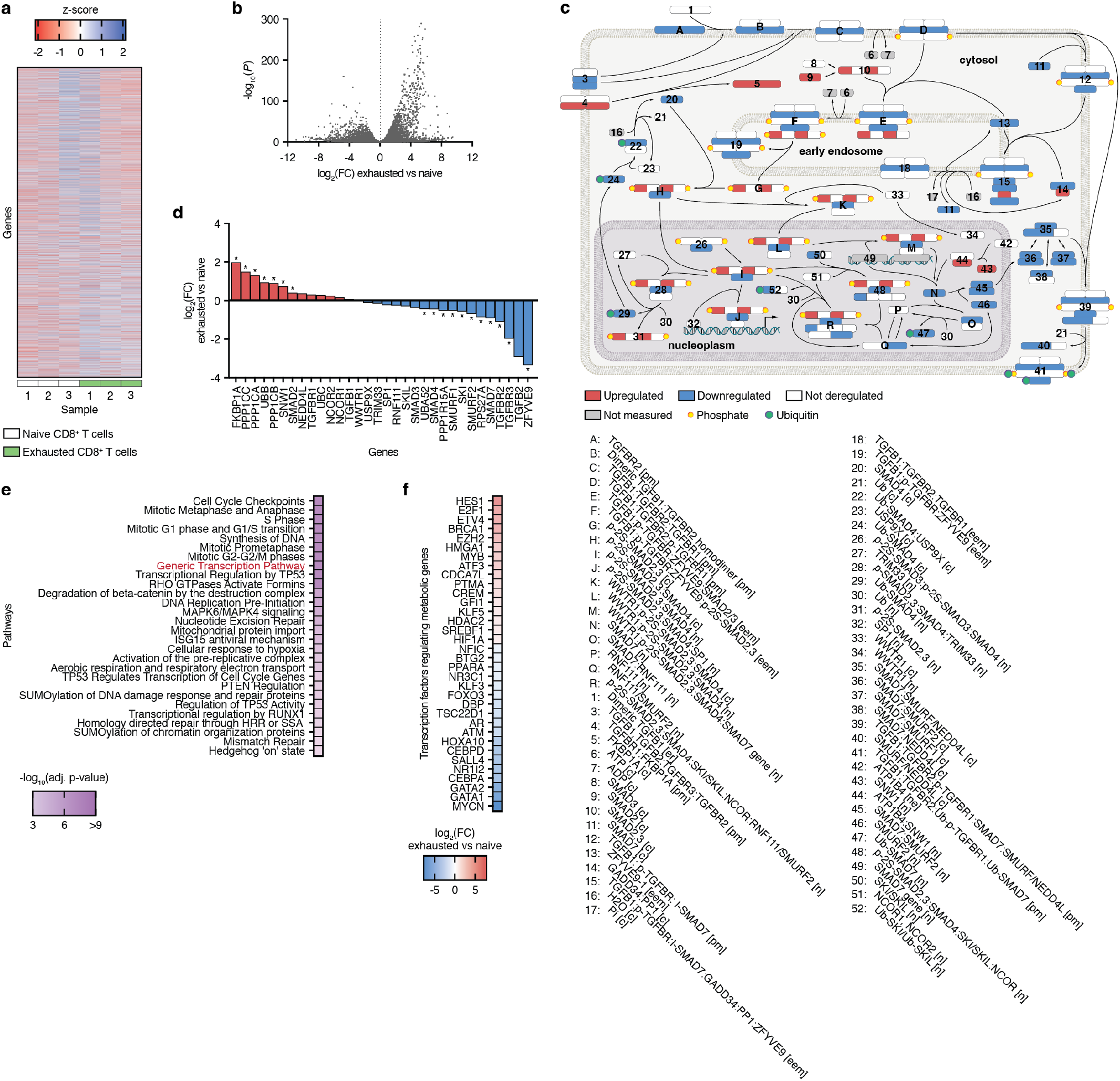
Differential expression between exhausted and naive CD8^+^ T cells and mapping onto TGF-β to SP1 cascades. **a**, Heatmap of normalized expression, z-scored by row, for naive (n = 3) and exhausted (n = 3) CD8^+^ T cell samples used in this study, publicly available in Gene Expression Omnibus (GEO) under accession number GSE227316. **b**, Volcano plot of differentially expressed genes between exhausted and naive CD8^+^ T cells. The y-axis represents −log_10_(adjusted *P* value), whereas the x-axis represents log_2_(fold change) in gene expression. Adjusted *P* values indicate significance of differential expression and were calculated with DESeq2 (Benjamini-Hochberg corrected). **c**, Mapping of differential expression between exhausted and naive CD8^+^ T cells onto the balanced cascade shared by the 17- and 20-step cores. Color indicates differential expression status for each UniProt-mapped gene represented in the cascade species (Methods). Protein-complex species show separate colors for the UniProt-mapped gene associated with each constituent protein. Upregulated genes (red): log_2_(fold change) ≥ log_2_(1.3) and adjusted *P* ≤ 0.05; downregulated genes (blue): log_2_(fold change) ≤ −log_2_(1.3) and adjusted *P* ≤ 0.05; others are not deregulated (white). **d**, Bar plot showing differential expression between exhausted and naive CD8^+^ T cells for all genes appearing across the TGF-β to SP1 alternative cascades. The y-axis shows log_2_(fold change) (exhausted vs naive). Positive log_2_(fold change) is shown in red; negative log_2_(fold change) in blue. Asterisks denote significance (adjusted *P* ≤ 0.05). **e**, Heatmap showing the most deregulated Reactome pathway annotations participating in the TF-specific network (adjusted p-value ≤ 0.001), calculated from differential expression. Enrichment was performed at the gene level with over-representation analysis using the upper tail of the hypergeometric cumulative distribution and Benjamini-Hochberg correction. Pathways present in the TGF-β to SP1 alternative cascades are labeled in red. **f**, Heatmap showing the fold change of the most deregulated TFs that regulate metabolic genes (|log_2_(fold change)| ≥ 1 and adjusted *P* ≤ 0.05), calculated from differential expression.

## Notes

https://github.com/EPFL-LCSB/sigma

### Competing Interest Statement

The authors have declared no competing interest.

