## Supplementary Information for "Mechanistic reconstruction of receptor-to-transcription factor signaling integrating prior knowledge and omics"

### Supplementary Figures

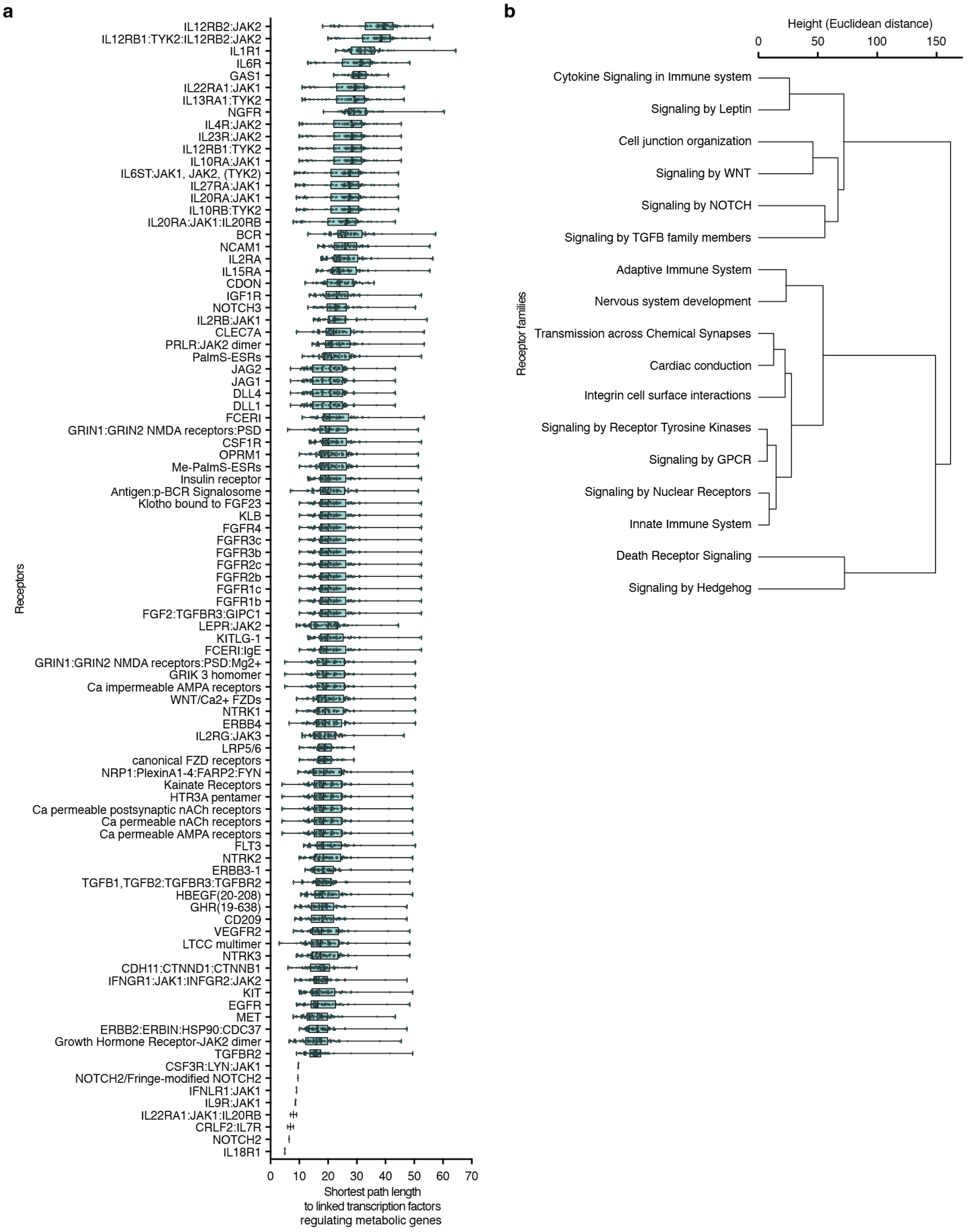

**Supplementary Fig. 1: Receptor and receptor-family path-length profiles to linked transcription factors.**

**a,** Box plots of shortest-path lengths from receptors to linked TFs regulating metabolic genes, ordered by the median per receptor. Each box spans the first to third quartiles with the median as a horizontal line. Whiskers indicate the minimum and maximum. Each point corresponds to a single TF. **b,** Hierarchical clustering of receptor families based on the average shortest-path length from receptors in each family to linked TFs regulating metabolic genes. Euclidean distance and Ward’s linkage were used.

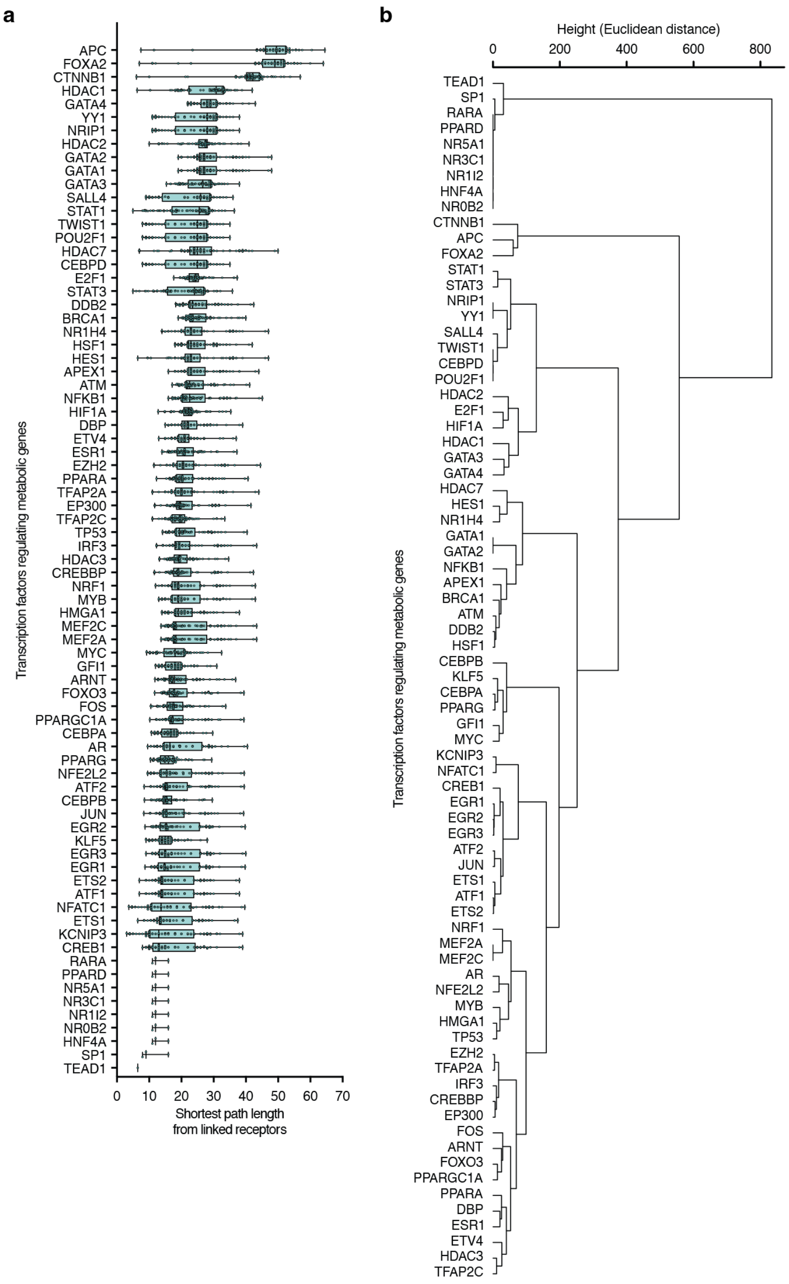


**Supplementary Fig. 2: Transcription factor path-length profiles from linked receptors.**

**a,** Box plots of shortest-path lengths from linked receptors to TFs regulating metabolic genes, ordered by the median per TF. Each box spans the first to third quartiles with the median as a horizontal line. Whiskers indicate the minimum and maximum. Each point corresponds to a single receptor. **b,** Hierarchical clustering of TFs regulating metabolic genes based on shortest-path lengths from linked receptors. Euclidean distance and Ward's linkage were used.

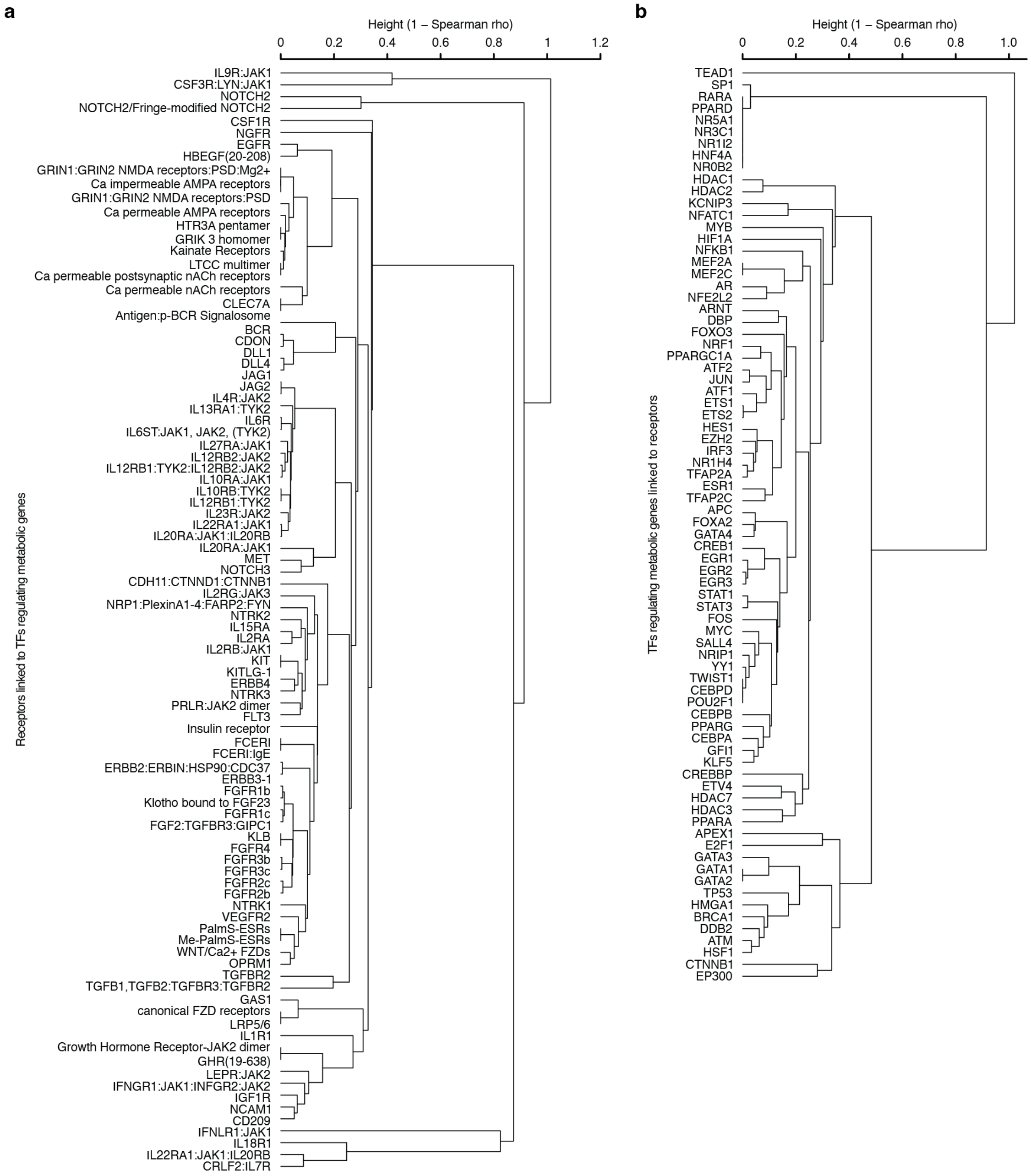


**Supplementary Fig. 3: Similarity of receptor-downstream and TF-upstream subnetworks.**

**a,** Hierarchical clustering of balanced subnetworks downstream of receptors linked to TFs that regulate metabolic genes (n = 93). Spearman distance between receptor-downstream species-weight vectors (Methods) and average linkage were used. **b,** Hierarchical clustering of balanced subnetworks upstream of TFs that regulate metabolic genes and are linked to receptors (n = 77). Spearman distance between TF-upstream species-weight vectors (Methods) and average linkage were used.

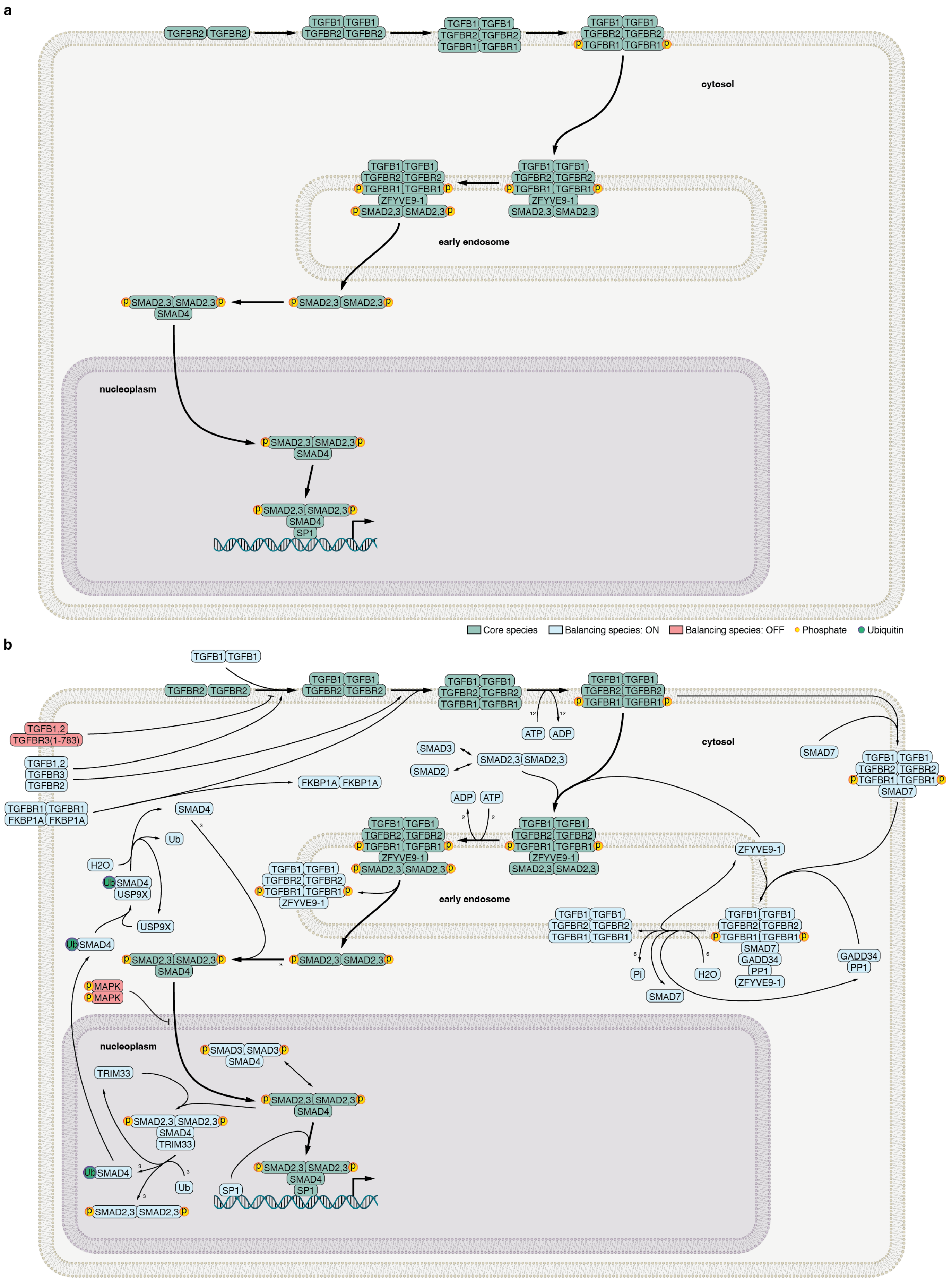
a
TGFBR2 TGFBR2
TGFB1 TGFB1
TGFBR2 TGFBR2
TGFBR1 TGFBR1
p TGFBR1 TGFBR1 p
cytosol
ZFYVE9-1
SMAD2,3 SMAD2,3
p SMAD2,3 SMAD2,3 p
early endosome
SMAD4
nucleoplasm
SP1
Core species
Balancing species: ON
Balancing species: OFF
Phosphate
Ubiquitin
b
TGFB1,2
TGFBR3(1-783)
TGFB1,2
TGFBR3
TGFBR2
TGFBR1 TGFBR1
FKBP1A FKBP1A
ATP
ADP
SMAD3
SMAD2
SMAD7
H2O
Ub
Ub SMAD4
USP9X
Pi
GADD34
PP1
p MAPK
TRIM33
p SMAD3 SMAD3 p
SMAD4
TRIM33
Ub SMAD4
p SMAD2,3 SMAD2,3 p
12
2
3
6

**Supplementary Fig. 4: Detailed TGF-β to SP1 cascade for the nine-step alternative.**

**a,** Nine-step shortest core cascade from TGFBR2 to the nuclear complex of SP1 with the SMAD2/3:SMAD4 heterotrimer. Directed edges indicate the order of signaling events. **b,** Elementally balanced cascade for the nine-step core in panel **a**, assembled with the minimal supporting reactions and species (23 reactions; 42 species). Compartments indicate cytosol, early endosome, and nucleoplasm; core species are shown in green; balancing species are shown in blue; inhibitor OFF states are shown in red; post-translational modifications (phosphorylation and ubiquitination) are marked where present in both panels.

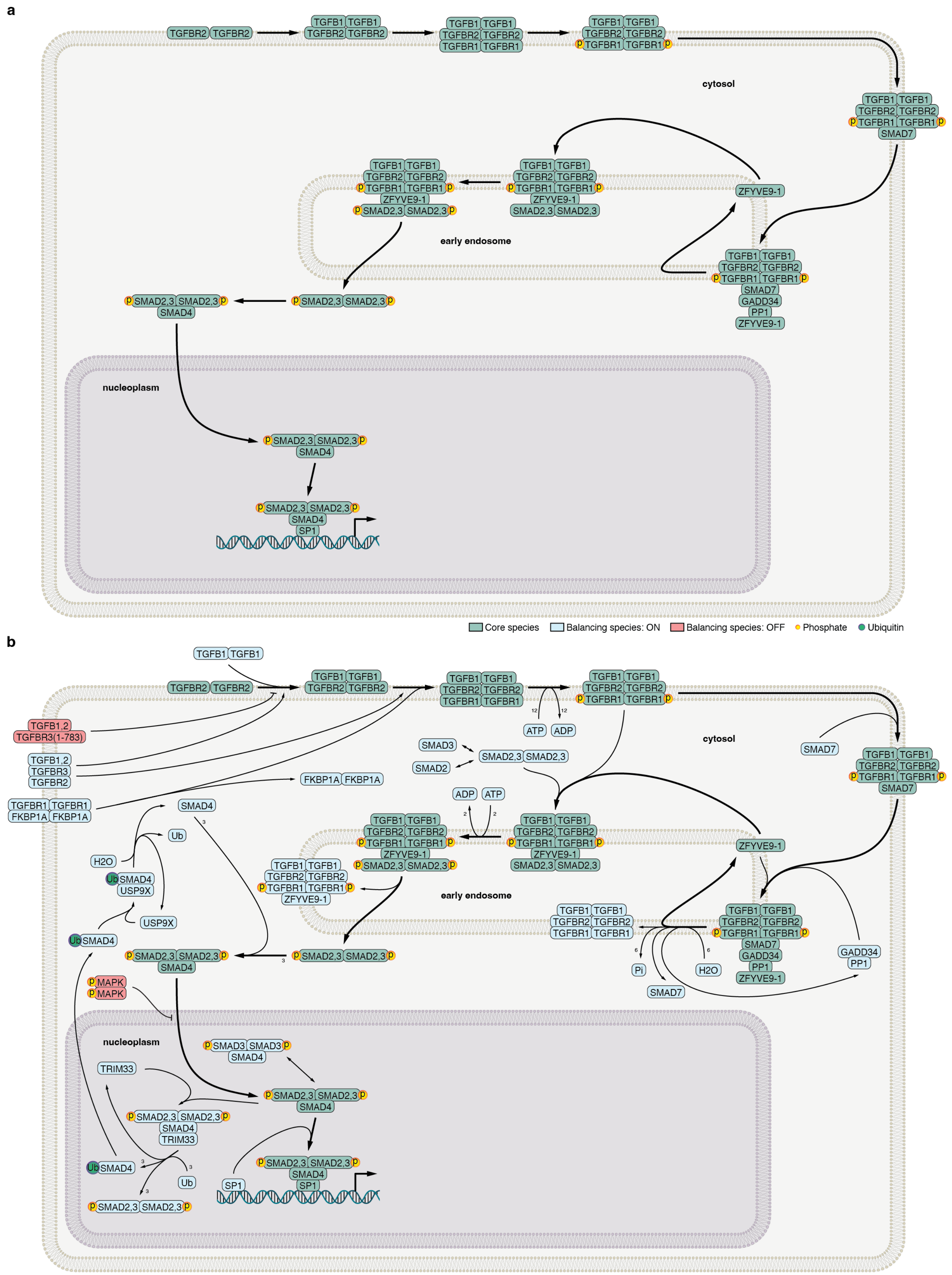
a
b
cytosol
early endosome
nucleoplasm
TGFBR2 TGFBR2
TGFB1 TGFB1
TGFBR1 TGFBR1
SMAD7
ZFYVE9-1
SMAD2,3 SMAD2,3
SMAD4
GADD34
PP1
SP1
Core species
Balancing species: ON
Balancing species: OFF
Phosphate
Ubiquitin
TGFB1,2
TGFBR3(1-783)
TGFBR3
FKBP1A FKBP1A
ATP
ADP
SMAD3
SMAD2
H2O
Ub
USP9X
MAPK
TRIM33
Pi
SMAD3 SMAD3

**Supplementary Fig. 5: Detailed TGF-β to SP1 cascade for the twelve-step alternative.**

**a,** Twelve-step alternative core cascade from TGFBR2 to the nuclear complex of SP1 with the SMAD2/3:SMAD4 heterotrimer. Compared with the nine-step core, this alternative branches via SMAD7 and SARA (ZFYVE9). Directed edges indicate the order of signaling events. **b,** Elementally balanced cascade for the twelve-step core in panel **a**, assembled with the minimal supporting reactions and species (23 reactions; 42 species). This core yielded the same elementally balanced cascade as the nine-step core. Compartments indicate cytosol, early endosome, and nucleoplasm; core species are shown in green; balancing species are shown in blue; inhibitor OFF states are shown in red; post-translational modifications (phosphorylation and ubiquitination) are marked where present in both panels.

a

b

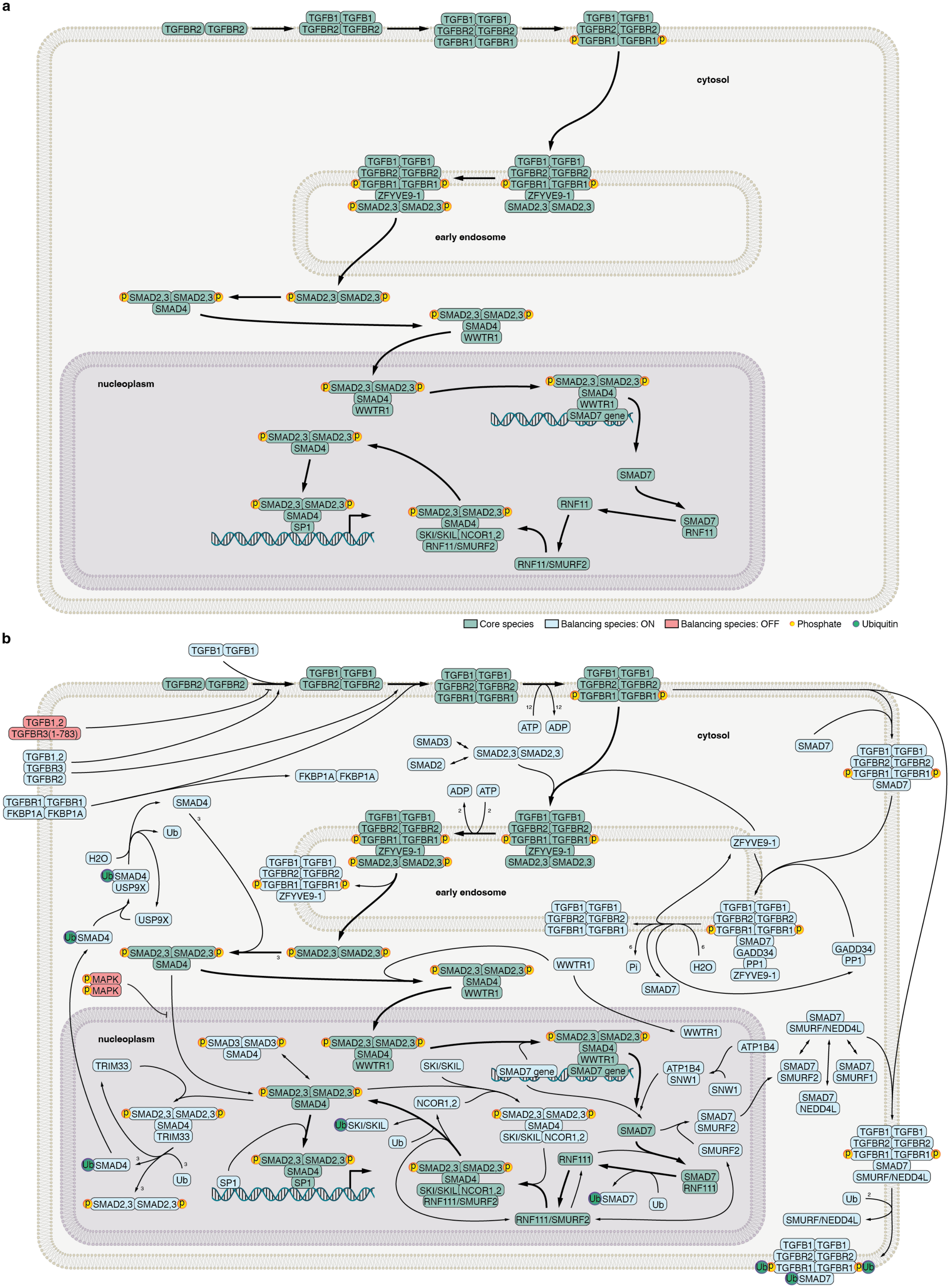

TGFBR2 TGFBR2
TGFB1 TGFB1
TGFBR1 TGFBR1
cytosol
ZFYVE9-1
SMAD2,3 SMAD2,3
early endosome
SMAD4
WWTR1
nucleoplasm
SMAD7 gene
SMAD7
RNF11
SP1
SKI/SKIL
NCOR1,2
RNF11/SMURF2
Core species
Balancing species: ON
Balancing species: OFF
Phosphate
Ubiquitin
TGFB1,2
TGFBR3(1-783)
TGFBR3
FKBP1A FKBP1A
ATP
ADP
SMAD3
SMAD2
H2O
Ub
USP9X
MAPK
Pi
GADD34
PP1
SMURF/NEDD4L
SMURF2
SMURF1
NEDD4L
ATP1B4
SNW1
TRIM33
SMAD3 SMAD3
RNF111
RNF111/SMURF2

**Supplementary Fig. 6: Detailed TGF-β to SP1 cascade for the seventeen-step alternative.**

**a,** Seventeen-step alternative core cascade from TGFBR2 to the nuclear complex of SP1 with the SMAD2/3:SMAD4 heterotrimer. Compared with the nine-step core, this alternative branches via WWTR1. Directed edges indicate the order of signaling events. **b,** Elementally balanced cascade for the seventeen-step core in panel **a**, assembled with the minimal supporting reactions and species (51 reactions; 70 species). Additional reactions and species are present owing to inclusion of the WWTR1-dependent branch relative to the balanced cascades of the nine- and twelve-step cores. Compartments indicate cytosol, early endosome, and nucleoplasm; core species are shown in green; balancing species are shown in blue; inhibitor OFF states are shown in red; post-translational modifications (phosphorylation and ubiquitination) are marked where present in both panels.

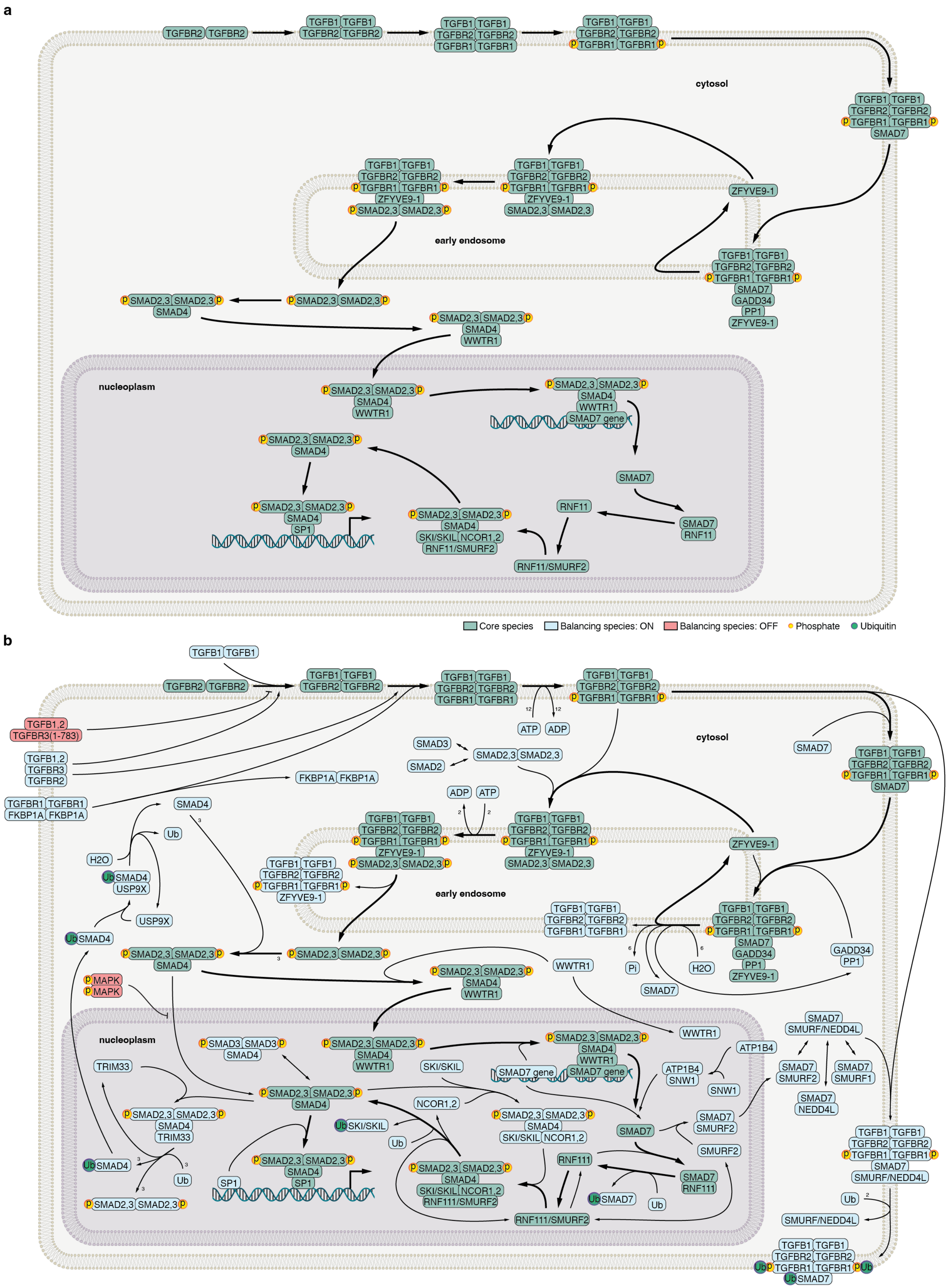
a
TGFBR2 TGFBR2
TGFB1 TGFB1
TGFBR1 TGFBR1
cytosol
SMAD7
ZFYVE9-1
SMAD2,3 SMAD2,3
early endosome
GADD34
PP1
SMAD4
WWTR1
nucleoplasm
SMAD7 gene
SP1
SKI/SKIL NCOR1,2
RNF11/SMURF2
RNF11
Core species
Balancing species: ON
Balancing species: OFF
Phosphate
Ubiquitin
b
TGFB1,2
TGFBR3(1-783)
TGFBR3
TGFBR2
FKBP1A FKBP1A
ATP
ADP
SMAD3
SMAD2
H2O
Ub
USP9X
MAPK
Pi
SMAD3 SMAD3
TRIM33
SKI/SKIL
NCOR1,2
RNF111
RNF111/SMURF2
SMURF/NEDD4L
SMURF2
SMURF1
NEDD4L
ATP1B4
SNW1

**Supplementary Fig. 7: Detailed TGF-β to SP1 cascade for the twenty-step alternative.**

**a,** Twenty-step alternative core cascade from TGFBR2 to the nuclear complex of SP1 with the SMAD2/3:SMAD4 heterotrimer. Compared with the seventeen-step core, this alternative branches via SMAD7 and SARA (ZFYVE9). Directed edges indicate the order of signaling events. **b,** Elementally balanced cascade for the twenty-step core in panel **a**, assembled with the minimal supporting reactions and species (51 reactions; 70 species). This core yielded the same elementally balanced cascade as the seventeen-step core. Compartments indicate cytosol, early endosome, and nucleoplasm; core species are shown in green; balancing species are shown in blue; inhibitor OFF states are shown in red; post-translational modifications (phosphorylation and ubiquitination) are marked where present in both panels.

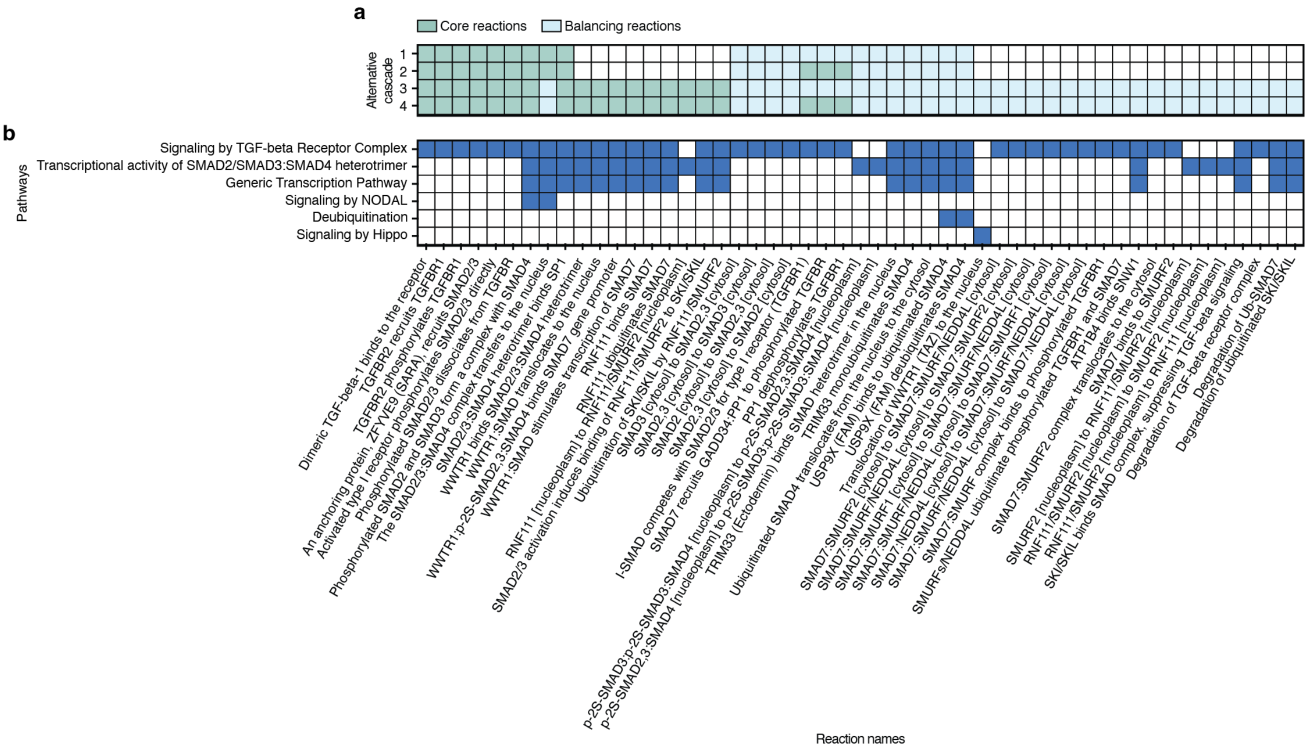


**Supplementary Fig. 8: Reaction participation and pathway membership in TGF-β to SP1 cascades.**

**a,** Summary map of reaction participation across alternative cascades. Columns list reaction names and are aligned with panel **b**. **b,** Mapping of reactions to pathway annotations. Colored boxes indicate Reactome pathway membership.

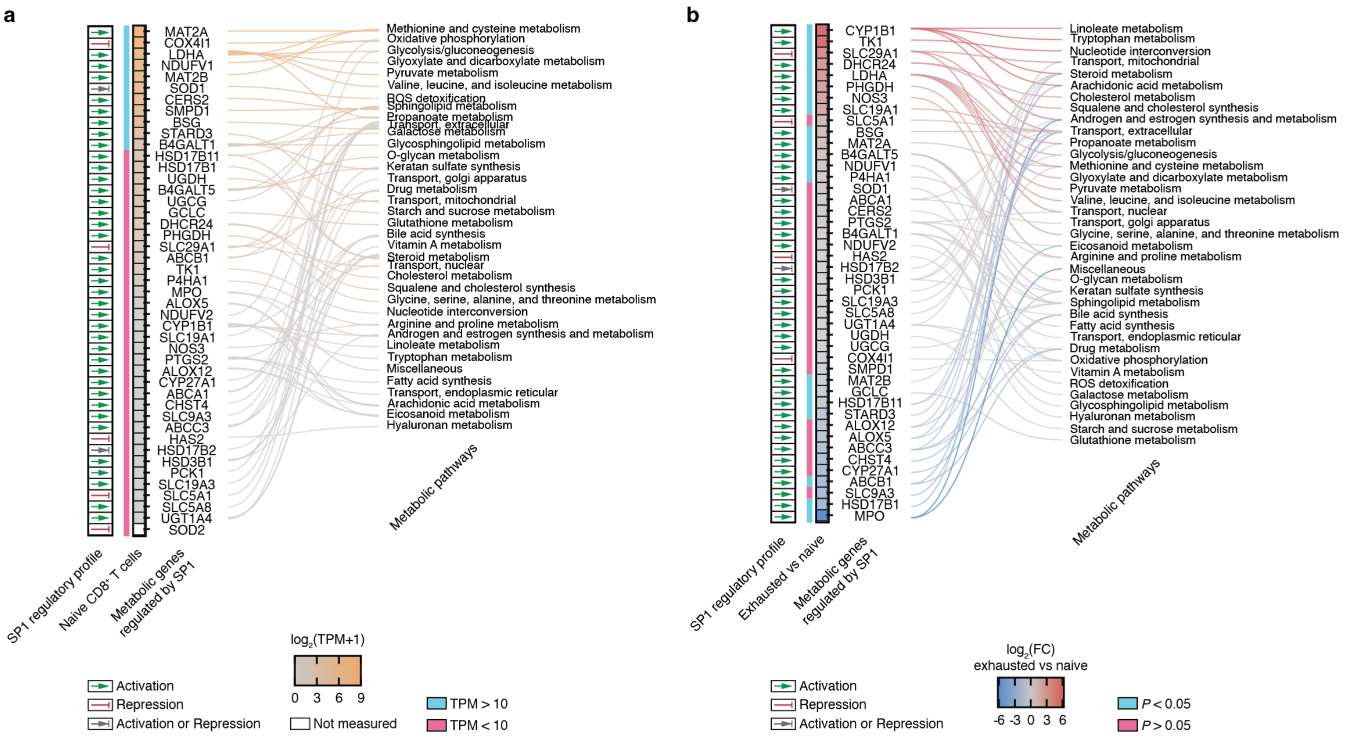


**Supplementary Fig. 9: Metabolic regulatory relationships of SP1 in naive and exhausted CD8$^+$ T cells.**

**a,** Mapping of absolute expression from naive CD8$^+$ T cells onto the regulatory relationships connecting SP1 to the metabolic genes it regulates and their associated metabolic pathways. The SP1 regulatory profile indicates whether each annotated regulatory relationship corresponds to activation or repression (based on TRRUST annotation). Heatmap shows gene expression in naive cells. TPM ≥ 10 (side band) marks highly expressed genes. **b,** Mapping of differential expression between exhausted and naive CD8$^+$ T cells onto the same SP1 regulatory relationships. Heatmap shows fold changes in exhausted versus naive cells. Adjusted $P$ values (side band) indicate significance of differential expression and were calculated with DESeq2 (Benjamini-Hochberg corrected).

### Supplementary Tables

#### Table S1: TF-specific signaling network reconstruction and characterization.

**a,** Regulatory relationships between transcription factors and metabolic genes. **b,** Metabolic subsystems regulated by transcription factor targets. **c,** Transcription factors regulating metabolic genes. **d,** Pathways in the TF-specific signaling network. **e,** Receptor-ligand interactions in the TF-specific signaling network. **f,** Pathway expansion across the TF-specific signaling network. **g,** Gap filling in the TF-specific signaling network. **h,** Cofactors and small molecules in the TF-specific signaling network. **i,** Effect of cofactors and small molecules on network topology.

#### Table S2: Receptor-to-TF connectivity in the TF-specific signaling network.

**a,** Receptor-to-TF connections. **b,** Connectivity of receptor families. **c,** Shortest paths from receptor families to transcription factors. **d,** Connectivity of transcription factor families. **e,** Species essentiality in receptor-to-transcription factor subnetworks. **f,** Correlations between receptor-downstream subnetworks. **g,** Correlations between TF-upstream subnetworks.

#### Table S3: TGF-β to SP1 cascade components and pathway annotations.

**a,** Species participation in TGF-β to SP1 cascades. **b,** Reaction participation in TGF-β to SP1 cascades. **c,** Gene mapping to species in TGF-β to SP1 cascades. **d,** Gene participation in TGF-β to SP1 alternative cascades. **e,** Pathway membership of species in TGF-β to SP1 cascades. **f,** Pathway membership of reactions in TGF-β to SP1 cascades. **g,** Fraction of TGF-β to SP1 alternative cascades assigned to Reactome pathways. **h,** Fraction of Reactome pathways covered by TGF-β to SP1 alternative cascades.

#### Table S4: Naive $CD8^+$ T cell expression mapping and TGF-β to SP1 cascade absolute enrichment.

**a,** Gene expression in naive $CD8^+$ T cells. **b,** UniProt-mapped expression in naive $CD8^+$ T cells. **c,** Expression mapped to TGF-β to SP1 cascades in naive $CD8^+$ T cells. **d,** Enrichment of TGF-β to SP1 alternative cascades in naive $CD8^+$ T cells. **e,** Pathway enrichment in naive $CD8^+$ T cells. **f,** Expression of transcription factors regulating metabolic genes in naive $CD8^+$ T cells. **g,** Expression of SP1-regulated metabolic genes in naive $CD8^+$ T cells.

#### Table S5: Exhausted versus naive $CD8^+$ T cell differential expression mapping and TGF-β to SP1 cascade differential enrichment.

**a,** Gene expression in naive and exhausted $CD8^+$ T cells. **b,** Differential expression in exhausted versus naive $CD8^+$ T cells. **c,** Differential expression mapped to TGF-β to SP1 cascades. **d,** Differential enrichment of TGF-β to SP1 alternative cascades. **e,** Pathway deregulation between exhausted and naive CD8+ T cells. **f,** Deregulation of transcription factors regulating metabolic genes. **g,** Deregulation of SP1-regulated metabolic genes.
